# Mitochondrial control of fuel switching via carnitine biosynthesis

**DOI:** 10.64898/2025.12.16.694762

**Authors:** Christopher Auger, Hiroshi Nishida, Bo Yuan, Guilherme Martins Silva, Masanori Fujimoto, Mark Li, Daisuke Katoh, Dandan Wang, Melia Granath-Panelo, Jihoon Shin, Rose Witte, Jin-Seon Yook, Anthony R.P. Verkerke, Alexander S. Banks, Sheng Hui, Lijun Sun, Shingo Kajimura

**Author notes:** Equal contribution (Hiroshi Nishida, Bo Yuan, Guilherme Martins Silva, Masanori Fujimoto).

## Abstract

Environmental adaptation often involves a shift in energy utilization toward mitochondrial fatty acid oxidation, which requires carnitine. Besides dietary sources of animal origin, carnitine biosynthesis from trimethyllysine (TML) is essential, particularly for those who consume plant-based diets; however, its molecular regulation and physiological role remain elusive. Here, we identify SLC25A45 as a mitochondrial TML carrier that controls carnitine biosynthesis and fuel switching. SLC25A45 deficiency decreased the carnitine pool and impaired mitochondrial fatty acid oxidation, shifting reliance to carbohydrate metabolism. *Slc25a45*-deficient mice were cold-intolerant and resistant to lipid mobilization by GLP1 receptor agonist (GLP-1RA), rendering them resistant to adipose tissue loss. Our study suggests that mitochondria serve as a regulatory checkpoint in fuel switching, with implications for metabolic adaptation and the efficacy of GLP-1RA-based anti-obesity therapy.

## INTRODUCTION

Eukaryotic cells possess an inherent ability to switch fuel sources in response to nutrient availability and physiological demands. This metabolic flexibility enables cells and tissues to utilize distinct pools of substrates, such as carbohydrates, lipids, and amino acids, in response to their energetic and biosynthetic needs (*1, 2*). This flexibility is particularly important during adaptation to fasting and cold exposure. During fasting, depletion of glycogen triggers a switch to fatty acid oxidation, derived from adipose tissue lipolysis, to fulfill energetic demands, while preserving glucose for the brain. Similarly, brown adipose tissue (BAT) undergoes a fuel shift from glucose to fatty acid and amino acid metabolism during cold adaptation, facilitating thermogenesis to maintain body temperature (*3–5*). Such flexibility in fuel utilization enhances metabolic efficiency and resilience, thereby supporting overall energy balance and metabolic health. Conversely, in pathological conditions such as heart failure, the heart shifts from predominantly fatty acid oxidation to an increased reliance on glucose metabolism, highlighting the versatility of fuel utilization (*6*).

An essential metabolite for fatty acid oxidation is carnitine. Carnitine facilitates the transport of long-chain fatty acids into the mitochondria, known as the carnitine shuttle, which involves three enzymes: carnitine palmitoyltransferase 1 (CPT1) for forming acylcarnitines in the outer mitochondrial membrane, carnitine-acylcarnitine translocase (CACT) for shuttling acylcarnitine across the inner membrane, and carnitine palmitoyltransferase 2 (CPT2) for liberating free fatty acids for β-oxidation in the mitochondrial matrix (*7*). While dietary sources of animal origin (for example, red meat) provide approximately 75% of total carnitine, de novo carnitine biosynthesis also contributes to the carnitine pool. Since most fruits and vegetables contain negligible amounts of carnitine, the majority of carnitine (>90%) is derived from de novo synthesis in strict vegetarians, vegans, and herbivores (*8, 9*). Carnitine deficiency leads to impaired fatty acid metabolism, resulting in hypoketotic hypoglycemia, muscle weakness, and cardiomyopathy, in which carnitine supplementation is an effective treatment to restore metabolic function (*9, 10*). Additionally, carnitine deficiency is associated with chronic kidney disease, which is accompanied by impaired removal of metabolic waste products, such as creatinine and blood urea nitrogen (*11, 12*).

De novo carnitine biosynthesis begins with the mitochondrial import of N6, N6, N6-trimethyllysine (TML), a precursor derived from dietary sources or protein degradation (*7*). Within the mitochondrial matrix, trimethyllysine hydroxylase, epsilon (TMLHE), catalyzes the conversion of TML to 3-hydroxy-6-N-trimethyllysine (HTML) and then undergoes sequential enzymatic transformations: 3-hydroxy-6-N-trimethyllysine aldolase (HTMLA) converts it to 4-N-trimethylaminobutyraldehyde (TMABA), which is further oxidized by 4-N-trimethylaminobutyraldehyde dehydrogenase (ALDH9A1) to form γ-butyrobetaine (γ-BB). Finally, γ-butyrobetaine dioxygenase (BBOX1), also known as γ-butyrobetaine hydroxylase (GBBH), catalyzes the synthesis of L-carnitine (**Fig. 1A**). This process requires the transport of TML across the inner mitochondrial membrane (IMM), which is impermeable to metabolites.

**Figure 1.**
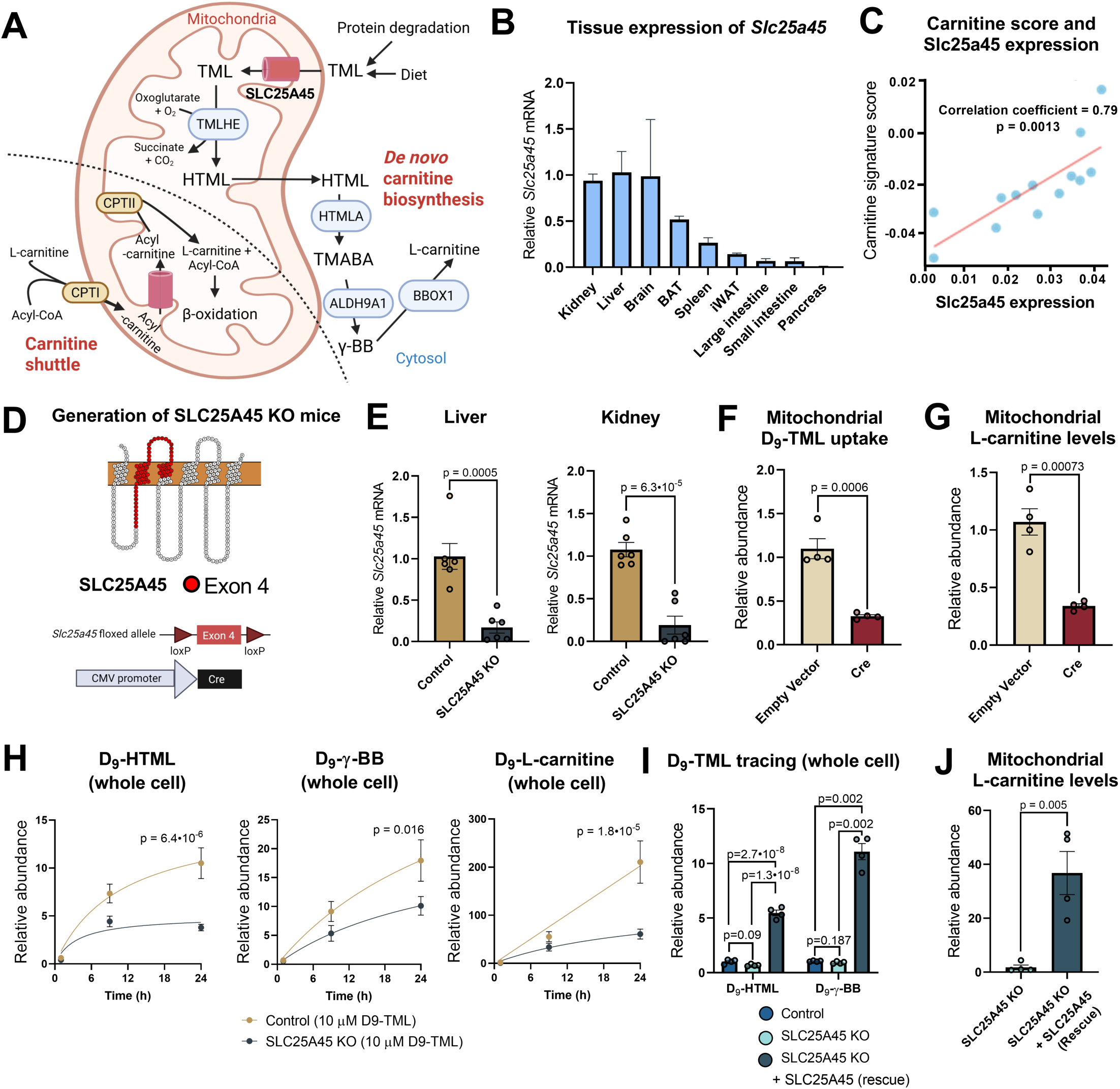
SLC25A45 is required for carnitine biosynthesis. **(A)** Schematic of the de novo carnitine biosynthesis pathway. **(B)** Tissue distribution of *Slc25a45* mRNA expression in the indicated mouse tissues, normalized to kidney expression (n = 3). **(C)** Correlation between *Slc25a45* expression and carnitine signature score. **(D)** Schematic of *Slc25a45* deletion by targeting Exon 4. **(E)** Relative mRNA expression of *Slc25a45* in the liver and kidney (n=6). (**F)** TML uptake in isolated mitochondria from control cells and SLC25A45 KO cells expressing Cre (n = 4). **(G)** Mitochondrial contents of carnitine in control and SLC25A45 KO cells in (F). **(H)** D_9_-TML tracing in whole-cell hepatocytes from control and SLC25A45 KO mice at the indicated time points. D_9_-HTML (left), D_9_-γ-BB (middle), and D_9_-carnitine (right) were quantified by LC-MS (n = 6). **(I)** D_9_-TML tracing in WT HEK293 cells, SLC25A45 KO, and KO cells expressing ectopic SLC25A45 (rescue) after 4 hours of incubation with D_9_-TML (n = 3). **(J)** Mitochondrial carnitine contents in SLC25A45 KO cells and rescue cells (n = 4). Bars represent mean and error, as shown in s.e.m. P values were calculated by Pearson correlation test (C), unpaired t-test [(E), (F), (G), and (J)], one-way ANOVA with Tukey’s post hoc HSD test (I), or two-way ANOVA with Šídák’s multiple comparisons test (H).

Notably, a genome-wide association study of metabolites (GWAS-METSIM) from 6136 Finnish participants (*13*) showed a strikingly significant correlation (*P* = 1 x 1.9^-80^) between TML and the *SLC25A45* gene locus (**Fig. S1A**). The Common Metabolic Diseases Knowledge Portal (https://hugeamp.org) (*14, 15*) lists the *SLC25A45* gene, including a rare coding variant (rs34400381; R285C) as having strong correlations with serum creatinine concentrations (HuGE score, 121800) and estimated glomerular filtration rate (eGRF) for creatinine (HuGE score, 350) (**Fig. S1B**). These data motivated us to test the hypothesis that SLC25A45 is involved in the mitochondrial transport of TML and L-carnitine synthesis *in vivo*.

## RESULTS

### SLC25A45 is required for de novo carnitine biosynthesis

SLC25A45 is expressed in several organs in mice, with the highest gene expression in the kidney, brain, and liver, all of which have high carnitine synthesis capacity from TML (*7, 16*), as well as the interscapular BAT, spleen, and inguinal white adipose tissue (WAT) (**Fig. 1B**). Intriguingly, SLC25A45 expression shows a significant (p=0.0013) positive correlation with the gene signature of carnitine metabolism (**Fig. 1C**) as well as the expression of individual genes, including *Tmlhe*, *Aldh9a1*, *Bbox1*, and *Cpt1a* (**Fig. S1C**). This relationship is selective to SLC25A45 because the carnitine synthesis pathway did not show correlation with SLC25A44, a mitochondrial BCAA carrier (*4*), or SLC25A47, a liver-specific mitochondrial carrier (*17*) (**Fig. S1D**).

To test the above hypothesis, we next developed *Slc25a45*^flox/flox^ mice targeting Exon 4 of the *Slc25a45* gene that encompasses two transmembrane regions (**Fig. 1D**). We then crossed these with *CMV*-Cre mice to generate whole-body *Slc25a45* null mice (*CMV*-Cre; *Slc25a44*^flox/flox^, herein SLC25A45 KO mice) (**Fig. 1E**). Additionally, we harvested and immortalized the stromal vascular fraction (SVF) from the inguinal WAT of *Slc25a45*^flox/flox^, from which we established SLC25A45 KO cells and control cells by infecting immortalized SVFs with a retroviral Cre (pMSCV-Cre) or empty vector as control (**Fig. S1E**). Subsequently, we harvested mitochondria from these cells in order to determine the mitochondrial uptake of D_9_-labeled TML. Metabolomics analysis by Liquid Chromatography-Mass Spectrometry (LC-MS) showed that mitochondrial uptake of D_9_-TML was significantly (p=0.0006) lower in SLC25A45 KO mitochondria than in control mitochondria (**Fig. 1F**). Similarly, D_9_-TML uptake into the mitochondria isolated from the liver of SLC25A45 KO mice was lower than that in the mitochondria of the control mouse liver (**Fig. S1F**). This reduction was accompanied by lower carnitine content in the SLC25A45 KO mitochondria than in controls (**Fig. 1G**), suggesting a reduced carnitine shuttle in the absence of SLC25A45.

Hepatocytes express all the enzymes required for L-carnitine synthesis from TML, including TMLHE, ALDH9A1, and BBOX1 (see **Fig. 1A**) (*7*). Thus, we next isolated primary hepatocytes from control and SLC25A45 KO mice and proceeded to perform whole-cell tracing using D_9_-labeled TML. Metabolomics analyses found that hepatocytes from wild-type control mice produced D_9_-TML-derived precursors of L-carnitine, namely HTML and γ-BB, as well as D_9_-carnitine in a time-dependent manner (**Fig. 1H**). In contrast, SLC25A45 KO hepatocytes produced smaller amounts of D_9_-HTML, γ-BB, and L-carnitine. These results suggest that SLC25A45 is required for mitochondrial TML uptake as well as de novo synthesis of TML-derived carnitine. We note, however, that alternative pathways exist, as SLC25A45 KO cells produced detectable amounts of carnitine, albeit at lower levels than those in the controls. This possibility is implicated by the data showing that the expression of *Slc25a29,* a mitochondrial carrier for basic amino acids such as lysine (*18*), was elevated in SLC25A45 KO livers compared to controls (**Fig. S2A**). Furthermore, SLC25A45 KO hepatocytes were able to generate carnitine when γ-BB, a downstream metabolite of TML, was supplemented as a tracer (**Fig. S2B**).

Conversely, we examined the extent to which ectopic expression of SLC25A45 in KO cells could restore the pathway. To this end, we deleted SLC25A45 in HEK293 cells using a CRISPR system in which the cDNA sequence of human SLC25A45 was reintroduced (KO + SLC25A45, herein rescue cells) (**Fig. S2C**). Subsequently, we performed whole-cell metabolite tracing in wild-type cells, KO cells, and rescue cells using D_9_-TML. Metabolomics analyses found that SLC25A45 rescue cells contained higher amounts of D_9_-labeled HTML and γ-BB than SLC25A45 KO cells (**Fig. 1I**). Note that HEK293 cells expressed low levels of endogenous SLC25A45 and thus, there was no statistically significant difference between wild-type cells and SLC25A45 KO cells. We also observed similar results when SLC25A45 was overexpressed in wild-type HEK293 cells (**Fig. S2D**). Mitochondrial carnitine contents were restored when SLC25A45 was reintroduced in KO cells (**Fig. 1J**). We should note here that a previous report suggested the possibility that SLC25A45 could transport β-nicotinamide mononucleotide (NMN) (*19*). However, mitochondrial uptake assays using tritium-labeled NMN as a tracer failed to detect significant differences in mitochondrial NMN levels among wild-type, SLC25A45 KO, and rescue cells, which suggested that it is unlikely to be a specific substrate for SLC25A45 (**Fig. S2E**). Together, these results suggest that SLC25A45 is required for mitochondrial TML uptake and subsequent synthesis of L-carnitine from TML. We note that these results are consistent with recently published work (*20, 21*), which shows that SLC25A45 is required for the mitochondrial uptake of methylated amino acids, such as dimethylarginine and trimethyllysine, as well as carnitine synthesis.

### Reconstitution of TML transport by SLC25A45

To reconstitute TML transport across the IMM via SLC25A45, we used an *E. coli* strain that effectively expresses IMM proteins in the bacterial inner membrane by fusing them in-frame to maltose-binding protein (MBP) for correct orientation (*22–24*). We constructed control *E. coli* expressing an empty vector and MBP-SLC25A45 expressing *E. coli* (**Fig. S3A**). Subsequently, these cells were incubated with D_9_-TML, quickly washed, and subjected to LC-MS to measure their transport activity. We found that D_9_-TML uptake in SLC25A45-expressing bacteria increased relative to vector controls in a time-dependent manner, peaking at 60 minutes of incubation (**Fig. 2A**). Next, we assessed the substrate specificity of SLC25A45 by examining whether it transports structurally similar substrates, such as γ-BB and choline. While we observed significantly (P=0.00013) higher TML transport in SLC25A45-expressing *E. coli* versus controls, we did not find significant uptake of either γ-BB or choline (**Fig. 2B**).

**Figure 2.**
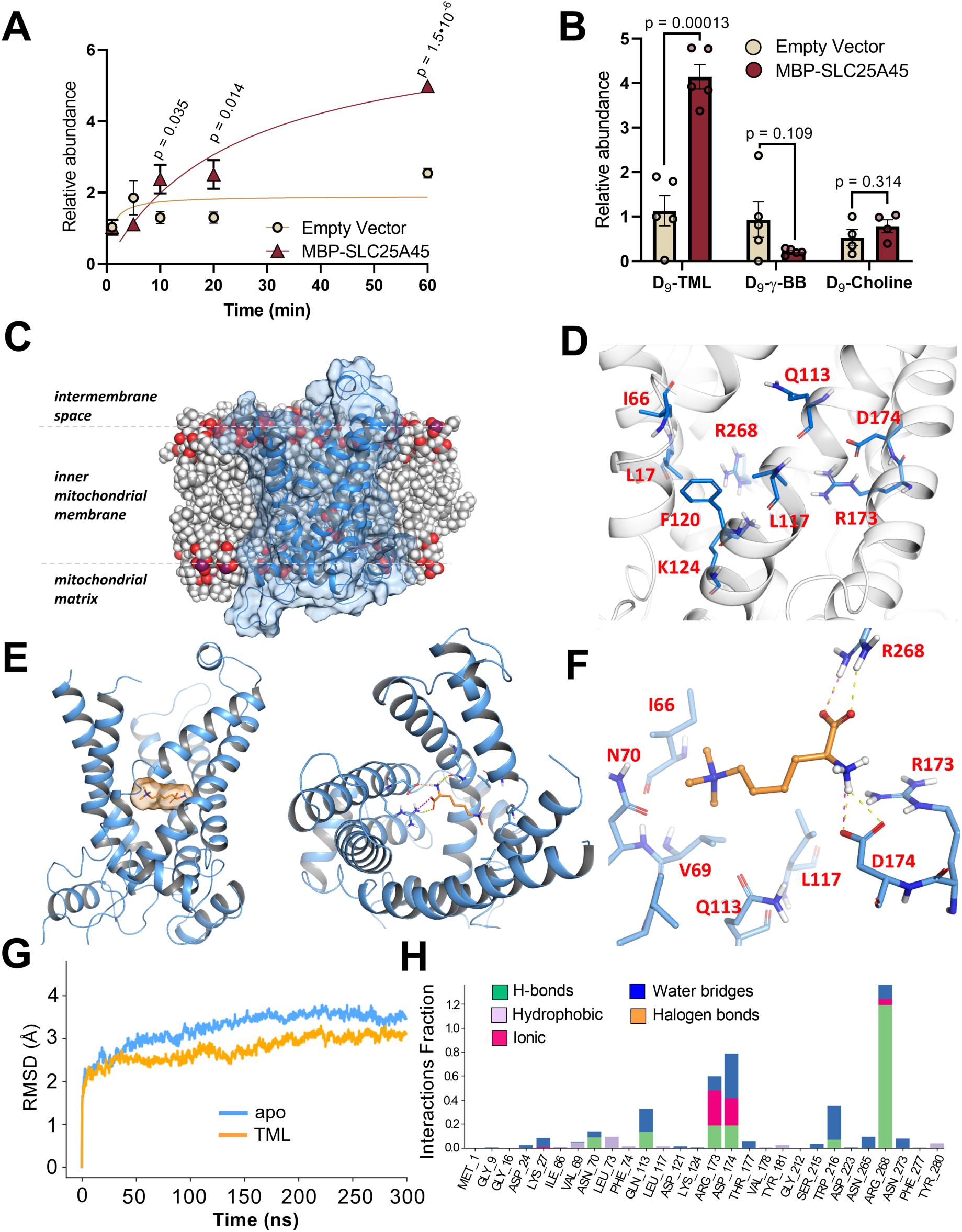
Reconstitution of SLC25A45-mediated TML transport. **(A)** Time-dependent D_9_-TML uptake in *E. coli* cells expressing an empty vector (control) and SLC25A45 fused in frame to MBP (MBP-SLC25A45) (n=4). **(B)** Substrate specificity of SLC25A45 in *E. coli* cells expressing MBP-SLC25A45 (n =5 for D_9_-TML and D_9_-γ-BB, n=4 for D_9_-choline). **(C)** Side view of SLC25A45 structure (AF-Q8N413-F1-v4; blue ribbons and surface) at the inner mitochondrial membrane. **(D)** Key amino acid residues (blue) at the central channel-like cavity. **(E)** Left: Side view of TML (orange carbons and surface) within the SLC25A45 binding cavity. Right: Top view of TML within the SLC25A45 binding cavity. **(F)** Close-up 3D view of TML (orange carbons in ball and stick) key interactions based on the MD replicate simulations. **(G)** Average RMSD against time plot obtained by MD simulations for SLC25A45 *apo* and complexed with TML. **(H)** Interaction bar chart for one of the TML-SLC25A45 MD replicate simulations. Bars represent mean and error, as shown in s.e.m. P values were calculated by two-way ANOVA with Šídák’s multiple comparisons test (A) or unpaired t-test (B).

To gain structural insights into how SLC25A45 selectively binds and transports TML as its substrate, we applied the following computational prediction models. Given the absence of experimentally resolved 3D structures for SLC25A45, we first used the AlphaFold (*25*) predicted structure for SLC25A45, which underwent a rigorous preparation protocol to ensure suitability for further methods. This predicted structure exhibits high-confidence scores and shares structural features with other known solute carriers (*26*), including a central channel-like cavity formed by six transmembrane alpha helices oriented perpendicular to the inner mitochondrial membrane (**Fig. S3B, C**). The structure represents the cytoplasmic-facing (c-state) conformation, in which the central cavity is open towards the cytosol (or intermembrane space), allowing potential substrates to access it (**Fig. 2C**). Notably, the channel’s interior surface is lined with the residues L17, I66, Q113, L117, F120, K124, R173, D174, and R268 (**Fig. 2D**). In sequence, we performed consensus docking followed by induced-fit docking protocols to predict the most energetically favorable and stable TML-SLC25A45 complex. We found that TML adopted a conformation that fit well within the SLC25A45 channel cavity (**Fig. 2E**). Specifically, this binding pose is primarily stabilized through specific interactions, including hydrogen bonds and/or salt bridges formed between the carboxylate of TML and the residues Q113, R173, and D174 of SLC25A45. In addition, hydrophobic contacts were observed between the *N*-trimethyl moiety of TML and the sidechains of residues L17, I66, V69, N70, L73, L117, F120, and K124 lining the cavity (**Fig. 2F, Fig. S3D, E**).

Next, we conducted multiple molecular dynamics (MD) simulations to investigate the behavior of SLC25A45 within the mitochondrial inner membrane, both in its *apo* state and in complex with TML. These simulations demonstrated that TML forms a stable and reproducible complex with SLC25A45. The root-mean-square deviation (RMSD) for the SLC25A45-TML complex averaged 2,74 Å, which was lower than the 3,23 Å observed for the apo state trajectories (**Fig. 2G**, **Fig. S4**). The average binding free energy (ΔG, calculated using the *MMGBSA* method) for the TML-SLC25A45 complex was -17.10 ± 3.75 kcal/mol, suggesting the binding of TML to SLC25A45 is energetically favorable (**Table S1**). In addition, we found that TML directed the carboxylate group towards R268 to establish a hydrogen bond, which was observed in approximately 90% of all simulations. Nonetheless, the *N*-trimethyl moiety kept close to the identical hydrophobic residues (**Fig. 2H, Fig. S5A**). This reproducible outcome suggests a subtle accommodation of TML within the channel center of the cavity. Furthermore, MD-based mutagenesis analysis revealed that a change in one of the key binding pocket amino acids (D174E) destabilizes the binding of TML to SLC25A45, reinforcing the high specificity of SLC25A45 to TML (**Fig. S5B**). SLC25A45 shares a structural homology to SLC25A48 that transports choline as a substrate (*27–29*), which also bears a *N*-trimethyl moiety (50% and 68% of sequence identity and conservation, respectively). Among the main residues surrounding the hydrophobic *N*-trimethyl moiety in the SLC25A45-TML complex, I66 and N70 are conserved (I69 and N73 in SLC25A48); however, L17, V69, L73, L117, and F120 are not conserved (V20, Y72, V76, G121, and V124 in SLC25A48) (**Fig. S5C**). These results align with experimental data that SLC25A45 transports TML but not choline.

### SLC25A45 loss results in systemic carnitine deficiency and a shift towards carbohydrate metabolism

To determine the biological significance of SLC25A45 in vivo, we examined the extent to which SLC25A45 controls systemic carnitine metabolism and fatty acid oxidation using SLC25A45 KO mice. Although we did not observe noticeable developmental defects in whole-body SLC25A45 KO mice, serum metabolomics identified lower amounts of L-carnitine and γ-BB in SLC25A45 KO mice compared to littermate control mice (**Fig. 3A**). In contrast, serum levels of L-carnitine precursors, L-lysine, and TML were higher in SLC25A45 KO than in control mice, which is in agreement with the results in cultured cells that SLC25A45 is required for the conversion from TML to γ-BB and L-carnitine. Carnitine levels in skeletal muscle, where more than 95% of the total carnitine is stored (*30*), as well as in the heart, were significantly (p=0.01) lower in SLC25A45 KO than in control mice (**Fig. 3B**). On the other hand, there was no difference in tissue carnitine contents in the BAT between the genotypes (**Fig. S6A**). We note that SLC25A45 KO mice exhibited a milder carnitine-deficient phenotype compared to that caused by primary carnitine deficiency, such as mutations in plasma membrane carnitine transporters, such as OCTN2. This may stem from the fact that SLC25A45 KO cells are obligated to generate carnitine from other sources (for example, from γ-BB), as the regular murine diet used in this study contains undetectable levels of carnitine (**Fig. S6B**).

**Figure 3.**
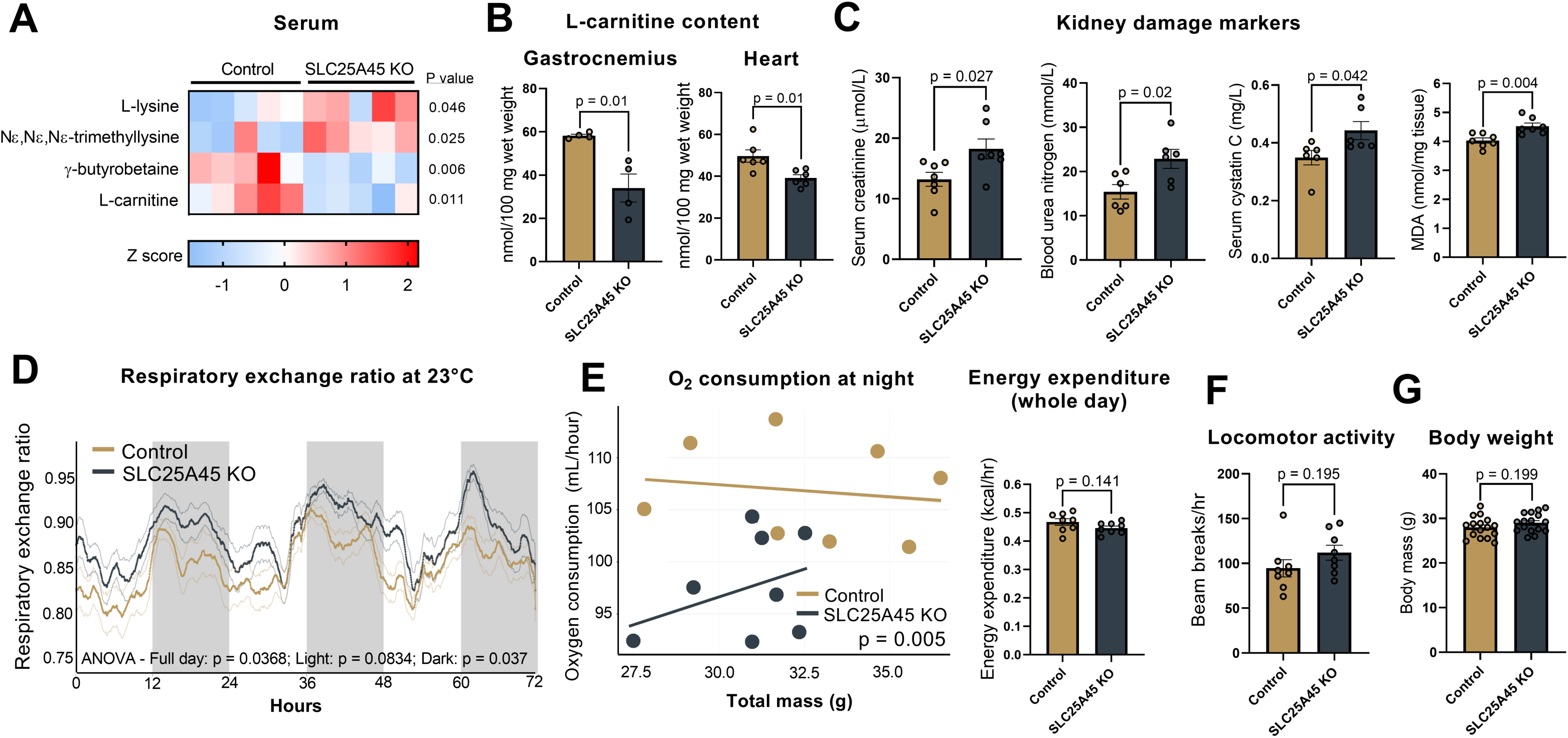
SLC25A45 loss results in systemic carnitine deficiency and altered fuel choice. **(A)** Serum levels of indicated metabolites from 3h fasted mice (n = 5). **(B)** Carnitine contents in the gastrocnemius muscle and heart after a six-hour fast (n=4 for quadriceps, n=6 for heart). **(C)** Serum levels of kidney damage markers (n=7 for serum creatinine and kidney MDA contents, n=6 for blood urea nitrogen and serum cystatin C). **(D)** Whole-body RER at room temperature (n=8). **(E)** Whole-body oxygen consumption rate and energy expenditure on a regular diet at room temperature (n=8). Left: Regression-based analysis of VO_2_ against body mass during nighttime. Right: Whole-day energy expenditure throughout the measurement. **(F)** Locomotor activity in (D). **(G)** Body weight of mice at 30 weeks of age (n=16). Bars represent mean and error shown as s.e.m. P values were calculated by unpaired t-test [(A), (B), (C), (F), and (G)], two-way ANOVA with Šídák’s multiple comparisons test (D), or CalR-ANCOVA with VO_2_ as a dependent variable and body mass as a covariate (E).

Systemic carnitine deficiency is associated with several pathogenic conditions, including chronic kidney disease with elevated circulating concentrations of creatinine and urea nitrogen (*11, 12*). This possibility is suggested by the human genetic association between *SLC25A45* and serum creatinine levels (*31*). In agreement, we found that several kidney damage markers, including serum creatinine concentrations, serum Cystatin C concentrations, and blood urea nitrogen concentrations, were higher in SLC25A45 KO mice compared to their littermate control mice (**Fig. 3C**). In addition, tissue concentrations of malondialdehyde (MDA), a marker of oxidative stress, were elevated in the kidneys of SLC25A45 KO mice relative to control mice. Histologically, the proximal tubules in the kidney of SLC25A45 KO mice displayed mild cytoplasmic vacuolation and pyknosis of the nuclei (**Fig. S7A**). Of note, SLC25A45 is highly expressed in the mouse kidney, where single-nuclei RNA-seq data (*32*) identified that epithelial cells of the proximal tubules and collecting ducts express *Slc25a45* and carnitine synthesis enzymes, including *Tmlhe* and *Aldh9a1* (**Fig. S7B**). Thus, we next developed kidney-specific SLC25A45 KO mice by crossing *Slc25a45*^flox/flox^ mice with *Ksp1.3*-Cre mice (also known as *Cdh16*-Cre, expressed predominantly in renal tubules) (*33*). However, there was no difference in the kidney damage markers and circulating carnitine concentrations between kidney-specific SLC25A45 KO mice and littermate control mice (**Fig. S7C, D**). As we discuss later, we also developed liver-specific SLC25A45 KO mice using *Albumin*-Cre mice, neither of which showed any changes in tissue content of carnitine and precursors (**Fig. S7E**). These data are in alignment with the model that carnitine is systematically derived from multiple organs (*8, 9*), and thus, tissue-specific deletion does not result in systemic carnitine deficiency. Accordingly, we used whole-body SLC25A45 KO mice for our in vivo physiology studies.

Next, we used indirect calorimetry to assess the extent to which SLC25A45 deletion altered fuel choice at the organismal level. The analyses showed that the respiratory exchange ratio (RER) of SLC25A45 KO mice was higher than that of littermate control mice, particularly during the dark phase at room temperature (23°C) (**Fig. 3D, Fig. S7F**). On the other hand, SLC25A45 KO mice displayed modest but significantly (p=0.005) lower VO_2_ levels at night compared to control mice, but not during the daytime (**Fig. 3E**). Thus, there was no significant difference in overall energy expenditure throughout the day between the genotypes. We also found no difference in locomotor activity (**Fig. 3F**), suggesting that altered RER and VO_2_ in SLC25A45 KO mice were not caused by immobility and behavioral changes. Accordingly, there were no significant differences in average body weight and organ weight between the genotypes on a regular chow diet (**Fig. 3G, Fig. S7G**).

Carnitine deficiency can result in hypoglycemia, as impaired fatty acid oxidation forces the body to rely more heavily on carbohydrate metabolism. Supporting this, blood glucose concentrations in SLC25A45 KO mice were significantly (p=0.005) lower than in littermate control mice following 12 hours of fasting (**Fig. 4A**). Furthermore, the blood concentrations of the ketone body beta-hydroxybutyrate (BHB) in SLC25A45 KO mice were lower than those in control mice after 24 hours of fasting (**Fig. 4B**). Additionally, SLC25A45 KO mice displayed higher glucose clearance than control mice on a standard chow diet (**Fig. 4C**). This phenotype was unrelated to altered insulin sensitivity because there was no difference in systemic insulin tolerance and fasting insulin concentrations between control and SLC25A45 KO mice (**Fig. S8A, B**).

**Figure 4.**
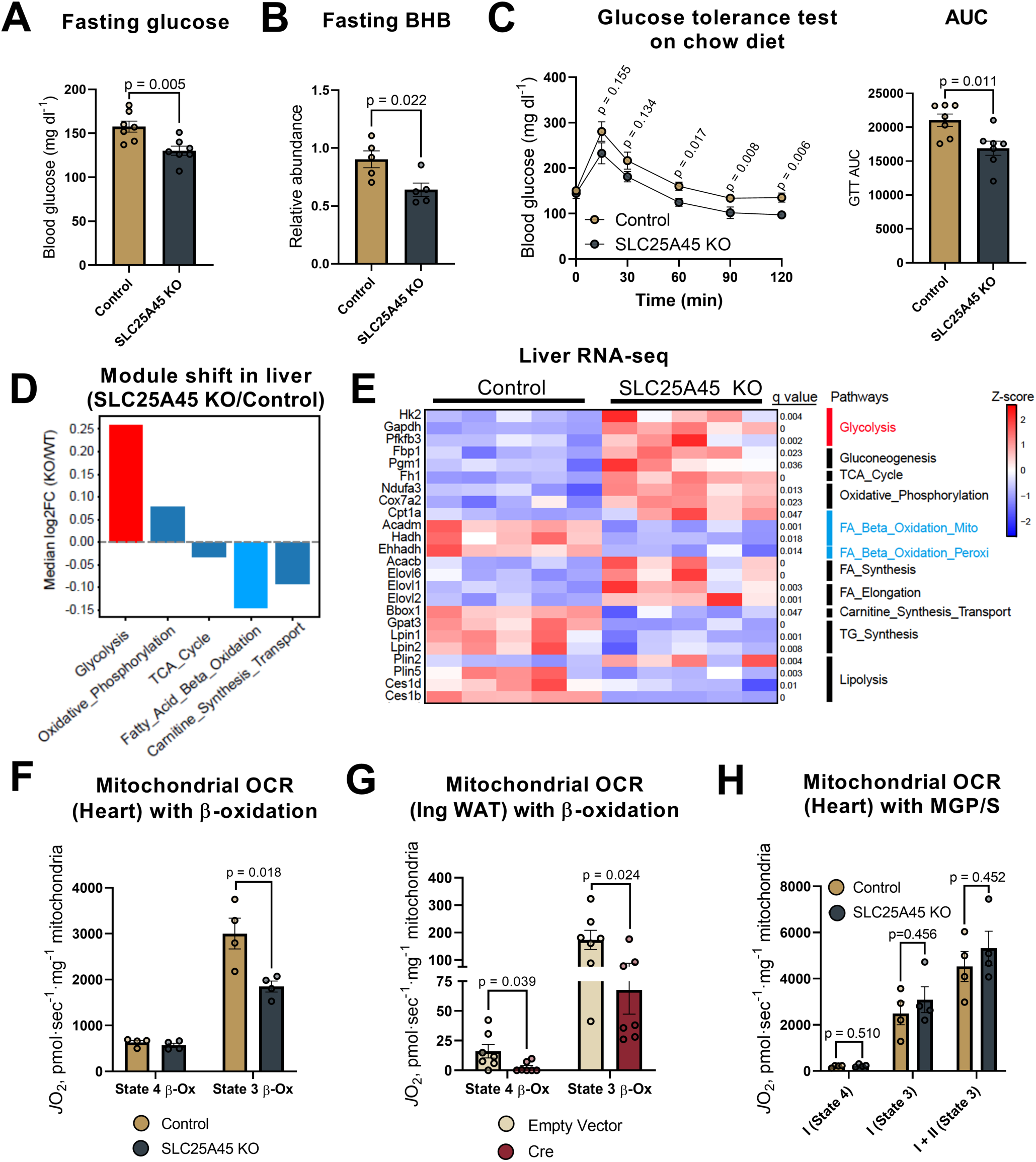
SLC25A45 deletion induces a fuel shift from fatty acid oxidation to carbohydrate/glucose catabolism. **(A)** Blood glucose levels on a standard diet (n=7). **(B)** Blood BHB levels after 24 hours of fasting (n=5). **(C)** Glucose tolerance test on a standard chow diet after 6 hours of fasting. Area of the curve (AOC) shown on the right (n=7). **(D)** The KEGG pathway analysis of the liver transcriptomics after 5 hours of fasting (n=5). **(E)** Z-score heatmap of indicated gene expression from (D). Pathways for each gene are listed on the right. **(F)** Mitochondrial State 3 and State 4 respiration (*J*O_2_) in the heart in the presence of palmitoyl-CoA, carnitine, and malate (n=4). **(G)** Mitochondrial respiration in inguinal WAT-derived adipocytes with palmitoyl-CoA, carnitine, and malate (n=7). **(H)** Mitochondrial respiration at indicated states was measured in the presence of glutamate, malate, and pyruvate (Complex I) and with ADP (State 3). Complex II-driven respiration was fueled with succinate (n=4). Bars represent mean and error shown as s.e.m. P values were calculated by unpaired t-test [(A), (B), (C), (F), (G), and (H)] or using the Benjamini-Hochberg procedure in R (E).

To examine the molecular changes in mice lacking *Slc25a45*, we next performed RNA-sequencing in the livers of SLC25A45 KO mice and littermate control mice. The transcriptomics identified 1675 genes that were altered between control and SLC25A45 KO mice (1078 genes upregulated, 597 genes downregulated). Among these changes, the KEGG pathway analysis showed that genes involved in the fatty acid beta-oxidation (for example, *Acadm*, *Hadh,* and *Ehhadh*) and carnitine synthesis (for example, *Bbox1*) were down-regulated in SLC25A45 KO mice relative to control mice, whereas the glycolysis (for example, *Hk2*, *Gapgh*, *Pfkfb3*) and mitochondrial OXPHOS pathways (for example, *Ndufa3, Cox7a2, Cpt1a*) were upregulated in SLC25A45 KO mice (**Fig. 4D**). In addition, genes involved in TG synthesis and lipolysis (for example, *Gpat3, Lpin1/2, Plin5*, and *Ces1d/b*) were downregulated in the livers of SLC25A45 KO mice, whereas fatty acid synthesis and elongation (for example, *Acacb, Elovl1,2,6*) were upregulated in the SLC25A45 KO livers (**Fig. 4E**). These results suggest that SLC25A45 loss in the liver induces a compensatory activation of glucose catabolism and fatty acid synthesis, while down-regulating mitochondrial fatty acid beta-oxidation. These changes were selective to the liver, as we found no significant changes in the glycolysis and beta-oxidation pathways in the heart (**Fig. S8C, D**).

To examine the functional consequences of impaired carnitine synthesis, we measured mitochondrial respiration in the heart, a primary site of long-chain fatty acid oxidation. We found that SLC25A45 KO mice had significantly (p=0.018) lower mitochondrial respiration than control mice at State 3 in the presence of ADP and palmitoyl-CoA (**Fig. 4F**). Similarly, we found lower mitochondrial respiration in SLC25A45 KO inguinal WAT-derived adipocytes compared to control cells when mitochondrial β-oxidation was assessed in the presence of palmitoyl-CoA (**Fig. 4G**). Decreased mitochondrial fatty acid oxidation of SLC25A45 KO mice is not a consequence of general mitochondrial dysfunction per se for the following reasons: First, SLC25A45 KO mitochondria exhibited equivalent oxygen consumption rate (OCR) to control mitochondria when pyruvate, glutamine, and succinate were provided (**Fig. 4H, Fig. S8E**). Second, when we added medium-chain fatty acid (C8, octanoate), which can access the mitochondrial matrix largely independently of the CPT1/CPT2-mediated carnitine shuttle in the kidney and liver (*34, 35*), the liver and kidney of SLC25A45 KO mice were able to increase OCR at an equivalent level to control mitochondria (**Fig. S8F)**. In addition, mitochondrial respiration was similar between the two groups in response to octanoylcarnitine. Third, mitochondrial morphology, as examined by electron microscopy, did not reveal any notable differences between SLC25A45 KO and control mice (**Fig. S8G**). Together, these results suggest that SLC25A45 deletion leads to a shift in fuel choice toward carbohydrates/glucose catabolism due to a reduced systemic carnitine pool and impaired long-chain fatty acid oxidation, while SLC25A45 KO mitochondria can utilize alternative fuels, such as pyruvate and medium-chain fatty acids.

### Carnitine biosynthesis via SLC25A45 is required for cold adaptation

Fuel switching to fatty acids is an essential adaptive process during cold exposure. We found that cold exposure elevated the expression of carnitine biosynthesis genes, including *Slc25a45*, *Tmlhe,* and *Bbox1* in the liver, BAT, and kidney, but not in the inguinal WAT (**Fig. 5A, Fig. S9A-C**). Furthermore, liver proteomics data (*36*) validated that SLC25A45 protein expression was elevated following cold exposure (**Fig. S9D**). These results showed that SLC25A45 is one of the cold-induced mitochondrial solute carriers.

**Figure 5.**
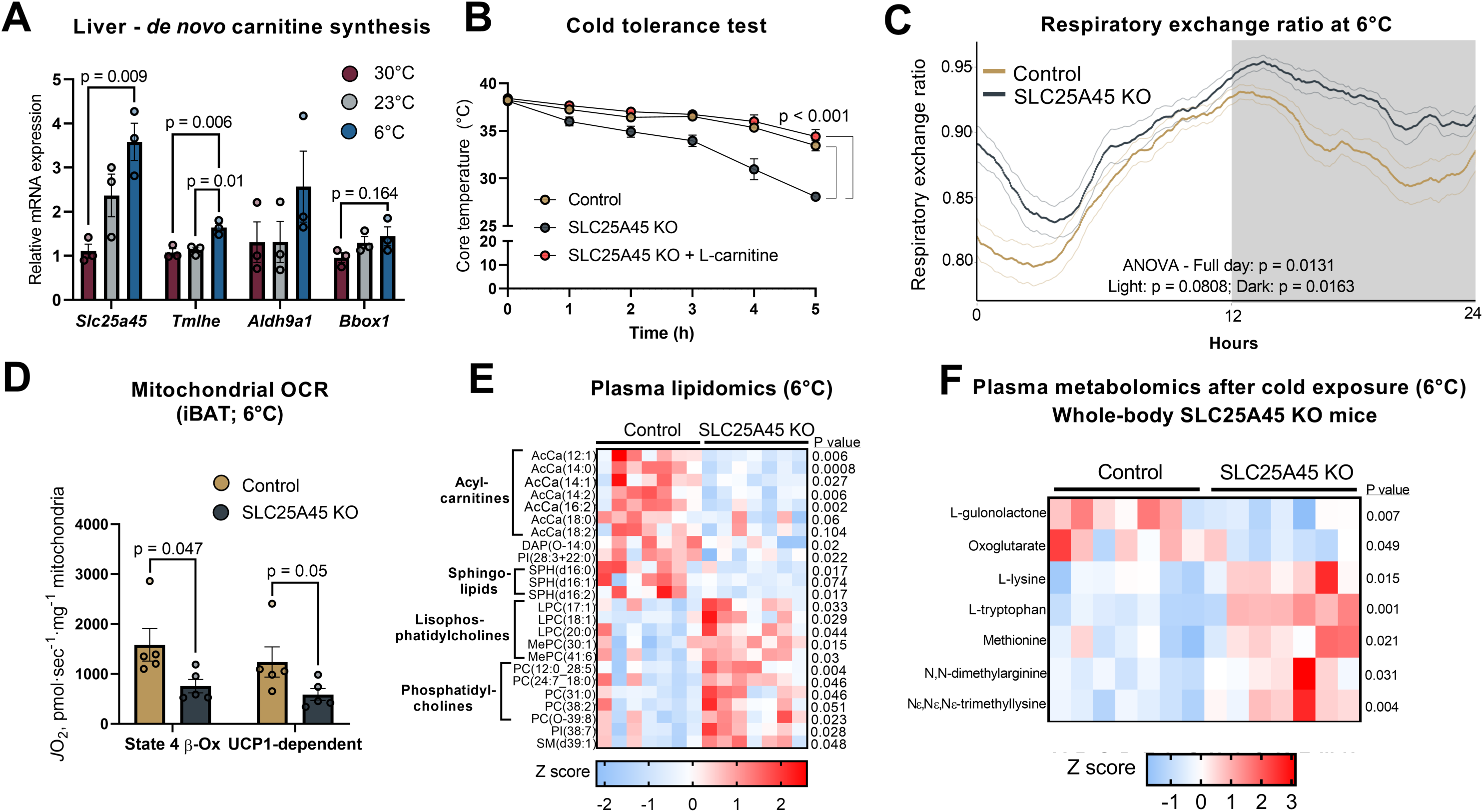
The carnitine biosynthesis pathway via SLC25A45 is required for cold adaptation. **(A)** mRNA expression of indicated genes in the liver of mice acclimated to 30°C, 23°C, and exposed to 6°C for 72 h (n = 3). **(B)** Cold tolerance test of standard diet-fed mice. Mice were housed at 30 °C for a week and placed in a cold chamber at 6 °C for the indicated time. A subset of SLC25A45 KO mice was supplemented with carnitine at 100 mg/kg for 5 days prior to cold exposure (n=5). **(C)** Whole-body RER during cold adaptation from 23°C to 6°C (n=8). **(D)** Mitochondrial *J*O_2_ in the BAT after cold exposure with palmitoylCoA, carnitine, and malate. UCP1-dependent respiration was determined in the presence of GDP (n=5). **(E)** Serum lipidomic after cold exposure (n=7). **(F)** Serum metabolomics in (E). Bars represent mean and error, as shown in s.e.m. Data in [(E) and (F)] are represented as z-scores. P values were calculated by one-way ANOVA with Tukey’s post hoc HSD test (A), Two-way ANOVA with Šídák’s multiple comparisons test [(B), (C)], or unpaired t-test [(D), (E), (F)].

The results led us to investigate whether SLC25A45-mediated carnitine synthesis is necessary for cold adaptation. When mice housed at 30°C were acutely exposed to cold temperature at 6°C, we found that the rectal temperature of SLC25A45 KO mice rapidly decreased and reached below 30°C after 5 hours of cold exposure (**Fig. 5B**). Conversely, carnitine supplementation (100 mg/kg for 5 days) prior to cold exposure completely rescued the hypothermic phenotype of SLC25A45 KO mice. Note that the mouse diets used in the study contain negligible amounts of carnitine (see Fig. S6B). The data suggest that cold intolerance in SLC25A45 KO mice is primarily due to systemic carnitine deficiency. During chronic cold acclimation from room temperature to cold at 6°C, the RER of SLC25A45 KO mice remained higher than that of control mice throughout the adaptation phase (**Fig. 5C**). To determine if the cold intolerance phenotype is caused by impaired BAT thermogenesis, we next isolated BAT mitochondria from SLC25A45 KO and littermate control mice following cold exposure, in which we measured mitochondrial respiration at State 4 in the presence and absence of GDP to assess UCP1-dependent respiration. We found that BAT-derived mitochondrial respiration in SLC25A45 KO mice was significantly (p≤0.05) lower than in controls under both conditions (**Fig. 5D**).

The liver is the major source of endogenous carnitine and expresses high amounts of SLC25A45 (see Fig. 1B), although other organs, such as the kidney, can generate carnitine (*8, 9*). Thus, we developed liver-specific SLC25A45 KO mice by crossing *Slc25a45*^flox/flox^ mice with *Albumin*-Cre mice (*Alb*-Cre; *Slc25a44*^flox/flox^, herein liver-SLC25A45 KO mice) (**Fig. S10A**). However, liver-SLC25A45 KO mice were able to maintain body temperature following cold exposure (**Fig. S10B**). In contrast to whole-body KO, liver-specific SLC25A45 deletion did not alter serum concentrations of γ-BB and carnitine (**Fig. S10C**). The results prompted us to examine circulating metabolites in SLC25A45 KO mice and liver-SLC25A45 KO mice. Lipidomics analysis identified that long-chain acylcarnitines, including C12:1-, C14:0-,14:1-, 14:2-, 16:2-, 18:0-carnitines, and several sphingolipids, were lower in the serum of whole-body SLC25A45 KO mice than in control mice following cold acclimation to 6°C (**Fig. 5E, Fig. S10D**). The result is consistent with a prior study showing that serum concentrations of acylcarnitines increased at 3 hours of cold exposure and thereafter in mice, where they served as key fuel for BAT thermogenesis and cold tolerance (*37*). We also found higher concentrations of lysophosphatidylcholines (LPC), phosphatidylcholines (PC), and sphingomyelin (SM) in the serum of SLC25A45 KO mice compared to control mice. Furthermore, serum metabolomics showed accumulated concentrations of L-carnitine precursor metabolites, including L-lysine, methionine, and TML, in SLC25A45 KO mice compared to control mice (**Fig. 5F**). On the other hand, there was no difference in circulating acylcarnitines, LPC, and PC concentrations between liver-SLC25A45 KO mice and littermate controls (**Fig. S10E**). These results underscore the importance of systemic carnitine availability in cold tolerance and circulating lipid profiles, which also explain why liver-specific deletion of SLC25A45 does not alter cold sensitivity in vivo.

### Carnitine biosynthesis is required for the optimal weight loss effect of GLP1 receptor agonists

We asked to what extent SLC25A45-mediated carnitine synthesis contributes to adaptive responses beyond cold acclimation. In this regard, food restriction is a good example of a fuel switch to fatty acids for ensuring sustained ATP synthesis while preserving glucose for the brain. Such a metabolic response can be pharmacologically induced by GLP1 receptor agonists (GLP-1RA) in mice with ad libitum access to food. GLP-1RA treatment promotes fatty acid oxidation with lowering RER in rodents and humans (*38, 39*). Accordingly, we examined the degree to which SLC25A45 loss affects the efficacy of GLP-1RA on whole-body energy balance. To test this, we administered a GLP-1RA, semaglutide, at a dose of 30 nmol/kg to SLC25A45 KO mice and littermate control male mice on a regular diet for 5 days (**Fig. 6A**). During the treatment, we found that SLC25A45 KO mice maintained higher RER than control mice (**Fig. 6B**). Even though the RER of both control and SLC25A45 KO mice was temporarily reduced by semaglutide treatment likely due to carnitines being synthesized through alternative routes, the RER in SLC25A45 KO mice remained higher than that in control mice right after semaglutide administration. On the other hand, there was no significant difference in VO_2_ and locomotor activities (**Fig. S11A, B**). Semaglutide injection led to elevated serum concentrations of long-chain acylcarnitines, particularly C12:0, 14:1, 14:2, 18:2, in control mice; however, such changes were not seen in SLC25A45 KO mice. Semaglutide also reduced serum triglycerides (TG) and diglycerides (DG) in control mice, whereas these lipids remained high in SLC25A45 KO mice. Additionally, there were differences in concentrations of some lysophosphatidylcholines (LPC) and phosphatidylcholines (PC) between the genotypes (**Fig. 6C**). These changes, like in cold exposure, are indicative of an impaired ability to mobilize lipid species in SLC25A45 KO mice. In addition, serum carnitine concentrations in SLC25A45 KO mice remained lower than those of control mice (**Fig. S11C**). These results further support the notion that SLC25A45 is required for fuel switching toward fatty acid oxidation in response to GLP-1RA-induced food restriction.

**Figure 6.**
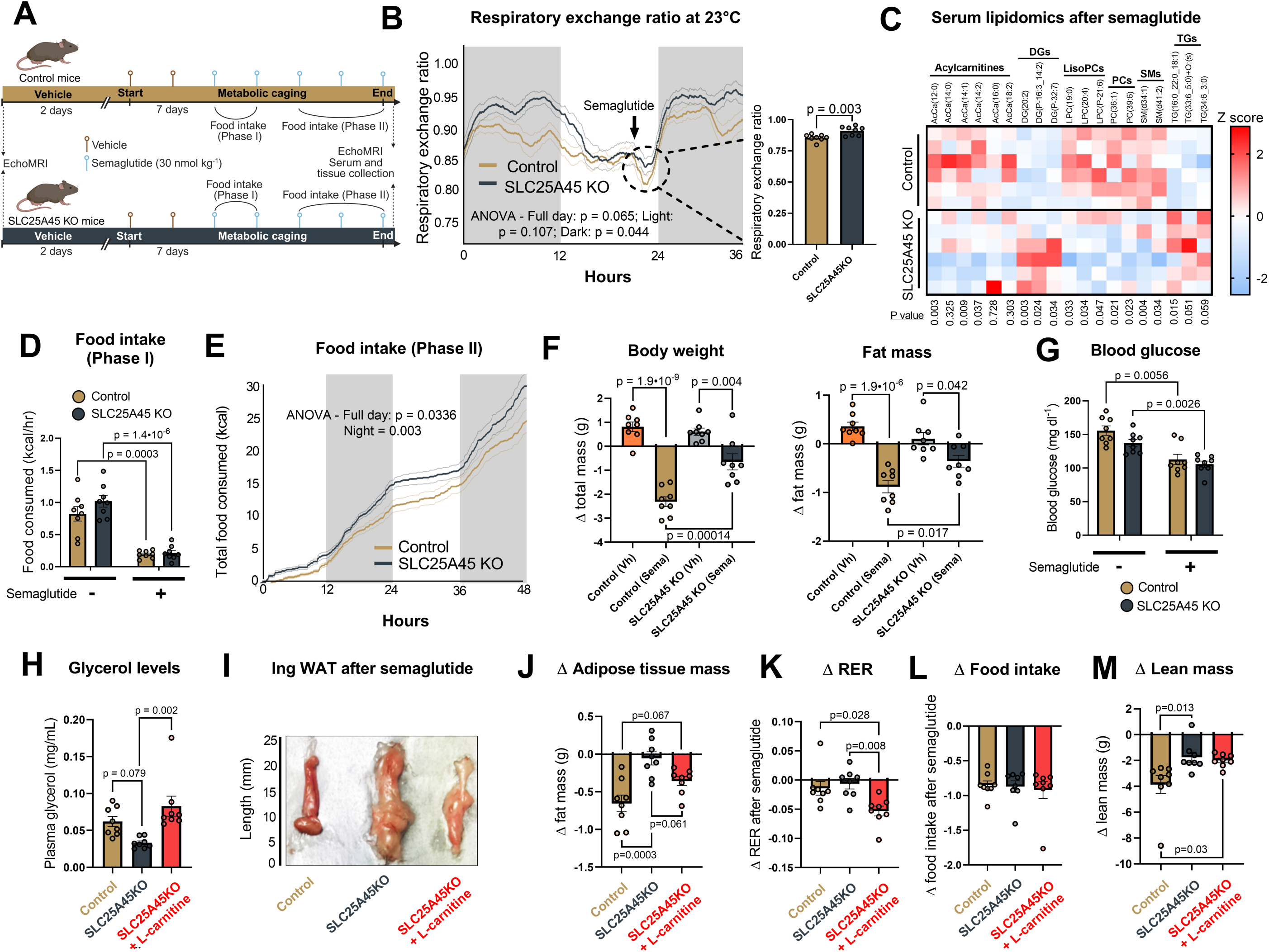
The carnitine synthesis pathway is required for the optimal anti-obesity effect of GLP-1RA. **(A)** Schematic of GLP-1RA experiment. Male mice on a standard chow diet were treated with semaglutide at 30 nmol/kg for 5 days. Energy expenditure, food intake, and locomotor activity were measured by indirect calorimetry. **(B)** Whole-body RER in response to semaglutide. Arrows indicate the time of semaglutide administration. The inset shows semaglutide-induced changes in RER (n=8). **(C)** Serum lipidomic analysis after semaglutide treatment represented as z-scores (n=6). **(D)** Food intake during Phase I in (A) (n=8). **(E)** Accumulated food intake during Phase II in (A). **(F)** Change in body weight and fat mass as measured by Echo-MRI after treatment with vehicle (Vh) or semaglutide (Sema) for 5 days. **(G)** Blood glucose levels after 24 hours of a single semaglutide injection (n=8). **(H)** Plasma glycerol concentration in SLC25A45 KO mice, littermate controls, and SLC25A45KO mice with L-carnitine supplementation at 100 mg/kg i.p. (n=8). **(I)** Representative images of inguinal WAT at the end of semaglutide treatment. **(J)** Adipose tissue mass by Echo-MRI (n=8). **(K)** Changes in RER during 30 min post-injection of semaglutide (n=8). **(L)** Change in food intake after acute semaglutide administration (n=8). **(M)** Lean mass by Echo-MRI after semaglutide treatment for 5 days (n=8). Bars represent mean and error shown as s.e.m. P values were calculated by Two-way ANOVA with Šídák’s multiple comparisons test [(B), (E)]), unpaired t-test (C), paired t-test [(D), (G)], or one-way ANOVA with Tukey’s post hoc HSD test [(F), (H), (J), (K), (L), (M)].

Semaglutide potently suppressed food intake in both control and SLC25A45 KO mice for the first day of semaglutide treatment (Phase I, **Fig. 6D**). However, the food intake of SLC25A45 KO mice was gradually restored and remained higher than that of control mice, particularly at night, during the Phase II of 5-day-treatment with semaglutide (**Fig. 6E**). Moreover, SLC25A45 KO mice were resistant to body weight loss and reduction in adipose tissue in response to semaglutide treatment compared to littermate control mice (**Fig. 6F**). On the other hand, semaglutide reduced glucose concentrations in both control and SLC25A45 KO mice to a similar degree (**Fig. 6G**). The data suggest that the SLC25A45-mediated carnitine pathway is required for semaglutide’s anti-obesity action, but not for its glucose-lowering effect.

Considering the critical role of lipolysis for fatty acid oxidation during fasting, we then tested if the carnitine biosynthesis pathway via SLC25A45 is required for adipose tissue lipolysis following GLP-1RA treatment. We did not observe a significant difference in plasma glycerol concentrations in SLC25A45 KO mice compared to control mice after GLP-1RA treatment, but carnitine supplementation restored adipose tissue lipolysis of SLC25A45 KO mice (**Fig. 6H**). Consistent with these changes, SLC25A45 KO mice were resistant to GLP-1RA-induced adipose tissue loss, whereas carnitine supplementation during the GLP-1RA treatment was effective in restoring the effect of GLP-1RA on adipose tissue loss in SLC25A45 KO mice (**Fig. 6I, J, and Fig. S11D**). Moreover, carnitine supplementation to SLC25A45 KO mice via intraperitoneal injection was effective in restoring RER concentrations when co-administered with GLP-1RA, albeit temporarily (**Fig. 6K**). This fast kinetics is consistent with the rapid turnover of carnitine in mice (*40*). These changes occurred independent of energy intake, as carnitine supplementation did not alter food intake during semaglutide treatment (**Fig. 6L**). Lastly, SLC25A45 KO mice retained higher lean mass than controls even after GLP-1RA treatment, although the difference persisted when SLC25A45 KO mice were supplemented with carnitine (**Fig. 6M**). Together, these results highlight the crucial role of carnitine biosynthesis via SLC25A45 in fuel switching from carbohydrates to fatty acids in response to cold acclimation, as well as GLP-1RA-induced body weight loss.

## DISCUSSION

The present study uncovers how mitochondria serve as a key regulatory checkpoint in fuel switching during adaptation. Specifically, we showed that SLC25A45 mediates the mitochondrial import of TML across the IMM, which is required for carnitine biosynthesis in vivo. Notably, a defect in the carnitine biosynthesis pathway by deletion of SLC25A45 results in impaired fatty acid oxidation in response to cold adaptation as well as GLP-1RA. The metabolic phenotype of SLC25A45 KO mice can be restored by carnitine supplementation, supporting the crucial role of SLC25A45 for carnitine biosynthesis. With these findings, an exciting question is to explore how SLC25A45-mediated carnitine synthesis contributes to a broader range of metabolic adaptations, including endurance exercise, intermittent fasting, and time-restricted feeding, as well as the response to fibroblast growth factor 21 (FGF21), the central hormone that coordinates fuel switching toward fatty acid oxidation (*41, 42*). In the meantime, these results also provide the following opportunities.

First, we need to understand how food-derived carnitine vs. de novo synthesis coordinately maintains the systemic carnitine pool. In humans, food-derived carnitines from meat, fish, and dairy account for approximately 75 % of the carnitine pool, while strict vegetarians and vegans largely (∼90%) depend on biosynthesis (*7, 10*). This suggests the existence of compensatory mechanisms to enhance carnitine biosynthesis in response to dietary carnitine availability (*30*). The identification of SLC25A45 as a gatekeeper of carnitine biosynthesis enables us to explore the regulatory mechanisms. We found that cold exposure induces SLC25A45 and several carnitine synthesis enzymes, including TMLHE and BBOX1, in the liver. Of note, thyroid hormone, which is a well-known activator of fatty acid oxidation and the carnitine shuttle (*43*), is reported to activate *SLC25A45* transcription in the liver via the binding of thyroid hormone receptor to the thyroid response element (TRE) on the promoter region of the *SLC25A45* gene (*43*). Consistently, thyroid hormone also promotes carnitine synthesis and fatty acid oxidation by activating transcription of the *BBOX* and *CPT1* genes (*44, 45*). An open question is to understand how the carnitine biosynthesis pathway, mediated by SLC25A45, is coordinately regulated by diet-derived carnitine and hormonal cues.

The second is related to the response to GLP-1RA. While recent advances in GLP-1RA therapy have revolutionized the treatment of obesity (*46–48*), emerging evidence highlights considerable individual variability in the efficacy of GLP-1RA-induced body weight loss (*49*). Although genetic variants in the GLP-1 receptor (GLP1R) gene have been shown to influence treatment response (*50–52*), what determines “responders” vs. “non-responders” remains poorly understood. A notable finding in the present study is that the carnitine synthesis pathway via SLC25A45 is required for the optimal body weight loss effect of the GLP-1RA; SLC25A45 KO mice exhibited elevated RER, impaired lipid mobilization, and decreased adipose tissue loss relative to control mice following GLP-1RA treatment. Carnitine supplementation to SLC25A45 KO mice was effective in restoring fatty acid oxidation and body weight loss effects of GLP-1RA. Moreover, the suppression of food intake by GLP-1RA was gradually reversed in SLC25A45 KO mice to a greater extent than in controls, indicating a compensatory drive to seek dietary carnitine. Future investigation is needed to better understand the mechanisms through which SLC25A45 loss leads to hyperphagia following chronic GLP-1RA treatment. Nonetheless, the present study raises the possibility that a total carnitine pool may influence the efficacy of GLP-1RA-mediated weight loss in individuals with limited dietary carnitine intake, such as vegans, who largely depend on the de novo synthesis pathway. Considering the reports that nutritional carnitine supplementation may be effective in body weight loss in some instances, but with mixed results (*53, 54*), it is conceivable that dietary carnitine supplementation may enhance the efficacy of GLP-1RAs for individuals with lower carnitine levels, including strict vegetarians and vegans.

## MATERIALS AND METHODS

### Mice

All procedures performed in this study were approved by the Institutional Animal Care and Use Committee (IACUC, protocol# 028-2022-25) at Beth Israel Deaconess Medical Center. Mice were housed at 23°C or 30°C (thermoneutral) in ventilated cages under a 12 h-12 h light/dark cycle. Mice were fed a standard diet (Lab Diet 5008) with free access to food and water unless specified. LabDiet 5008 is largely composed of long-chain (C16-18) fatty acids and TAGs. The diet lacks detectable carnitine (**Fig. S6B**). *Slc25a45^flox-/^*mice on a C57BL/6J background were generated in house using *Easi*-CRISPR with long single-stranded DNA as a donor to facilitate homology-directed repair (**Table S2**). LoxP sites were inserted on either side of exon 4 which codes for two transmembrane domains of SLC25A45. Whole body SLC25A45 KO mice were generated by crossing floxed mice with CMV-cre mice (B6.C-*^tg(CMV-cre)1Cgn^/*J, 006054). Liver SLC25A45 KO mice were generated by crossing floxed mice with *Albumin*-cre mice (B6.Cg-*Speer6-ps1^Tg(Alb-cre)21Mgn^*/J, 003574). Kidney SLC25A45 KO mice were generated by crossing floxed mice with *Ksp*-cre mice (B6.Cg-*^Tg(Cdh16-cre)91Igr^*/J, 012237).

### Cell culture

The stromal vascular fractions (SVFs) from inguinal WAT of male 8 week old *Slc25a45^flox-/^* mice were isolated by collagenase digestion per the established protocol (*55*) and immortalized by expressing the SV40 large T antigen as previously described (*56, 57*). Preadipocytes differentiation was induced by culturing cells with DMEM medium (with 10% FBS) and an adipogenic cocktail consisting of 5 mg/ml insulin, 1 nM T3, 0.5 mM rosiglitazone, 0.5 mM isobutylmethylxanthine, 125 nM indomethacin and 2 mg/ml dexamethasone. After two days, the cell medium was changed to one containing 10% FBS, 5 mg/ml insulin, 1 nM T3 and 0.5 mM rosiglitazone for another 5–7 days. HEK293T cells were cultured in DMEM (Gibco; 11965092) containing high glucose, 10% FBS and 1% penicillin–streptomycin.

For hepatocyte culture, primary cells from the liver of SLC25A45 KO and littermate control mice were isolated by the Liberase (Millipore Sigma; 5401119001) perfusion method. Cells were washed with hepatocyte wash medium (Thermo Fisher Scientific, Waltham, MA; 17704024), purified by 30% Percoll (GE Healthcare, Chicago, IL; 17089101) density gradient separation, and resuspended in William’s E medium (Thermo Fisher Scientific; 12551032) with 10% fetal bovine serum, 1X Glutamax, and 1% pen/strep. Cells were then seeded on collagen-coated plates at a final density of 3.5 × 10^4^ cells/cm.

### Plasmid and virus production

CRISPR/Cas9 plasmids for control and SLC25A45 KO in 293 cells (Santa Cruz Biotechnology; sc-418922, sc-410404, sc-410404-HDR) were transfected with UltraCruz transfection reagent and plasmid transfection medium (Santa Cruz Biotechnology; sc-395739, sc-108062). Retrovirus was produced in HEK293 packaging cells using 10 mg of plasmid and 5 mg each of the packaging plasmids VSV and gagpol via the calcium phosphate transfection method. The viral supernatant was collected and filtered (0.45 mm) 48 h later and 293 cells or preadipocytes were infected with virus and 10 mg/mL polybrene for 24 h. *Slc25a45^flox-/^* preadipocytes were infected with retrovirus expressing *cre* (34565, Addgene) or empty vector, followed by hygromycin selection at a dose of 200 μg ml^−1^. SLC25A45 KO 293 cells or WT 293 cells were infected with pMSCV-hSLC25A45 (VectorBuilder, VB220314-1244chs) and selected for with hygromycin (200 μg/ml) to generate stable rescue and SLC25A45 overexpression models.

### Mitochondrial D_9_-TML uptake assays

Confluent 15 cm dishes of differentiated SVF derived from Ing WAT were washed twice with ice-cold PBS then collected in ice-cold KPBS (136 mM KCl, 10 mM KH_2_PO_4_, pH 7.25). Cells were homogenized with 25 strokes of a Teflon glass homogenizer and centrifuged at 600g for 10 min at 4°C to remove cell debris. The supernatant was centrifuged at 9000g for 10 min at 4°C to obtain a mitochondrial pellet, which was resuspended in KBPS. Protein concentration was determined by BCA assay and 100 µg of mitochondrial protein was used for D_9_-TML uptake. Similarly, mitochondria were isolated from the livers of group-housed cold-exposed (6°C for 72h) control and SLC25A45 KO mice. Mitochondrial uptake buffer consisted of KPBS with 10 mM HEPES, 0.5 mM EGTA and 10 µM D_9_-TML. After 5 min at room temperature, mitochondria were pelleted and washed three times with ice-cold KPBS. Metabolites were extracted in 80% MeOH and subjected to MS.

### Whole-cell tracing

Primary hepatocytes from 16-week-old male mice were incubated with 10 µM D_9_-TML (MedChemExpress; hy-143711S) or 10 µM D_9_-γ-butyrobetaine (LGC Standards; TRC-B759497) for 1, 9, and 24 h in fresh William’s E medium with 10% fetal bovine serum, 1X Glutamax (Thermo Fisher Scientific; 30055061), and 1% pen/strep. 293T cells were incubated with 10 µM D_9_-TML for 4 h in standard growth medium (DMEM, 10% FBS, 1% P/S). After incubations, cells were washed with ice-cold PBS, scraped, and equal amounts of protein were used for metabolite extraction with ice-cold solvent (40% MeOH, 40% acetonitrile).

### Metabolomics

Chromatographic separation was achieved using a Vanquish UHPLC (Thermo Scientific, Waltham, MA, USA) and XBridge BEH Amide XP column (2.5 µm, 2.1 mm × 150 mm) with guard column (2.5 µm, 2.1 mm × 5 mm) (Waters, Milford, MA). Mobile phase A was water: acetonitrile 95:5, and mobile phase B was water: acetonitrile 20:80, with both phases containing 10 mM ammonium acetate and 10 mM ammonium hydroxide. The elution linear gradient was: 0 ∼ 3 min, 100% B; 3.2 ∼ 6.2 min, 90% B; 6.5. ∼ 10.5 min, 80% B; 10.7 ∼ 13.5 min, 70% B; 13.7 ∼ 16 min, 45% B; and 16.5 ∼ 22 min, 100% B, with flow rate of 0.3 mL/min. The autosampler was at 4°C. The injection volume was 5 µL. Needle wash was applied between samples using methanol: acetonitrile: water at 40: 40: 20. The mass spectrometry used was Q Exactive HF (Thermo Fisher Scientific, San Jose, CA), and scanned from 70 to 1000 m/z with switching polarity. The resolution was 120,000. Metabolites were identified based on accurate mass and retention time using an in-house library, and the isotopic labeling was analyzed by El-Maven.

### Reconstitution of TML uptake by SLC25A45

Bacterial expression of mitochondrial carriers and uptake assays was performed as described previously with modifications (*22–24*). MBP-tagged fusion hSLC25A45 was generated by combining the *E. coli* MBP containing the MalE signal peptide and the mitochondrial carrier gene. Each protein was combined with a short linker containing a thrombin cleavage site. MBP and hSLC25A45 coding sequences were synthesized as gBlock gene fragments (IDT) and cloned into the pET-21b(+) expression vector (NovoPro, V011022) with the In-fusion Cloning Kit (Takara, 638948). Expression of MBP-hSLC25A45 was performed in *E. coli* C43(DE3) cells (Biosearch Technologies, CMC0019-20X40UL). According to the manufacturer’s instructions, single aliquots of competent bacterial cells were transformed with pET21(+) containing MBP-hSLC25A45 and selected on LB agar plates containing 100 mg l^−1^ ampicillin (37 °C, overnight). Bacteria transformed with pET21 (+) empty or MBP-hSLC25A45 plasmid were cultured overnight in LB medium containing 100 mg/L ampicillin, followed by a 1:100 dilution in 300 ml of LB medium containing 100 mg/L ampicillin. Refreshed bacteria were incubated (37 °C, 200 r.p.m.) until OD600 reached 0.3-4, and then the cultures were removed and cooled on ice. Prepared bacteria were split into 50 ml aliquots in 250 ml flasks. Protein expression was induced by adding 0.1 mM isopropyl β-D-1-thiogalactopyranoside (IPTG, I6758-1G Sigma) to the chilled cultures, followed by induction overnight (20 °C, 300 r.p.m., 12-13 h).

To validate SLC25A45 protein expression in the bacterial system, homogenates from *E. coli* expressing empty vector or MBP-tagged SLC25A45 were separated on a 10% SDS-PAGE gel and transferred to PVDF membranes using the Trans-Blot Turbo Transfer System (Bio-Rad). PVDF membrane blots were blocked in Tris-buffered saline (TBS) with 0.1% Tween-20 and 5% BSA for 1 h and incubated overnight at 4C with mouse anti-MBP tag antibody (1:2000; Proteintech, 66003-1-IG; RRID:AB_11183040). Secondary antibody was anti-mouse IgG (1:5000, 31430; Thermo Fisher; RRID:AB_10960845). Induced cells were collected by centrifugation (4,000g, 15 min, 4 °C), washed once with ice-cold potassium HEPES buffer (10 mM HEPES + 50 mM KCl, pH 7.2), and centrifuged again (4,000g, 15 min, 4 °C). Bacteria were resuspended in ice-cold potassium HEPES buffer and OD600 was measured. Twenty OD600 mL units of bacteria transformed with empty vector or overexpressing MBP-hSLC25A45 were resuspended in 1 mL of potassium HEPES buffer. For timecourse assays, 10 µM D_9_-TML was added to the sample with shaking at room temperature and aliquots taken at 1, 5, 10, 20, and 60 minutes. For selectivity assays, 10 µM D_9_-TML, 10 µM D_9_-γ-butyrobetaine, or 10 µM D_9_-choline was added, and the reaction was stopped at 20 minutes. Samples were centrifuged at 10,000g for 5 min and washed twice with potassium HEPES buffer. Metabolites were extracted with ice-cold solvent (40% MeOH, 40% acetonitrile). MBP-hSLC25A45 fusion protein expression was assessed by Western Blots after the uptake assays.

### Structural Prediction of SLC25A45

The AlphaFold 2.0 structure was retrieved from AlphaFoldDB (https://alphafold.ebi.ac.uk/) under the code: AF-Q8N413-F1-v4. All figures were generated using *Pymol* 3.0.2. The AlphaFold structure was submitted to the *Protein Preparation Workflow* on *Maestro,* by *Schrodinger (Suite 2024-4)*, to reassign bond orders using CCD (Chemical Component Dictionary) database, add/replace hydrogens, fill in missing side chains, generate disulfide bonds, optimize hydrogen bonds with *PROPKA*, and perform restrained minimization to 0.30 Å root mean-square deviation (RMSD) using the *OPLS4* force field. Trimethyllysine (TML) canonical SMILES string was retrieved from PubChem (https://pubchem.ncbi.nlm.nih.gov/) – CAS 19253-88-4. *Ligprep* was used in default settings to convert the SMILES string to 3D structure and minimize it to a most energetically favorable conformation, considering *Epik* pH = 7.0 ± 2.0, up to 32 tautomer/stereoisomers, and *OPLS4* force field.

For docking studies, *Schrodinger’s* module *Receptor Grid Generation* was used to specify the ligand binding site based on the volume of the prioritized *Sitemap* pocket 1 within the SLC25A45 central channel. This grid corresponds to 20 and 10 Å of outer and inner enclosed boxes, respectively, and the centroid coordinates of x = 1.15, y = -0.23, and z = -1.17. This binding site was used to run TML consensus docking by using default parameters and up to 10 poses generation with GlideSP, GlideXP, AutodockVina, and JAMDA. The cluster with greatest number of poses was prioritized and all poses from it were rescored through the *Molecular Mechanics with Generalized Born and Surface Area (MMGBSA)* solvation module from *Schrodinger’s Prime* to estimate respective binding free energy ΔG values in kcal/mol. The best pose (lowest ΔG) was then used as reference for conducting *Induced-Fit Docking (IFD)*, by considering default settings and up to 20 poses generation per molecule. The resulting *IFD* poses were again rescored through *Prime MMGBSA* and clustered by visual inspection in groups with matching poses (orientation + conformation). We picked up the pose with the lowest ΔG free energy binding affinity value, in kcal/mol, to be elected as the most representative SLC25A45-TML docking complex.

### Molecular dynamics studies

Molecular Dynamics (MD) simulations were performed for the *apo* AlphaFold prepared structure of SLC25A45, as well as when in complex/docked to the ligand TML. *Desmond’s System Builder* was used to prepare each system comprising the *apo* protein or the protein-ligand complex. These were solvated with water TIP3P solvent model and neutralized with addition of ions and buffer NaCl at 0.15 M. The OPM webserver was used to obtain a mitochondrial inner membrane orientation model (POPC at 300 K) for the SLC25A45 prepared structures. Also, we considered orthorhombic boxes with minimized volumes, keeping each side distanced by a minimum distance of 10 Å from protein atoms. The force field OPLS4 was used and all the other options were set to default values. MD runs were conducted using *Desmond* by producing trajectories of 300 ns (*ca.* 1000 frames), 1.2 energy for recording interval, isothermal-isobaric (NPT) ensemble at 300 K and 1.01325 bar, and allowing relaxation of system before simulation. Each run was repeated 3 times (three replicates). Jobs were written and conducted on the O2 High Performance Compute Cluster, supported by the Research Computing Group, at Harvard Medical School, through our licensed software, part of the SBGrid Consortium. Produced trajectories were submitted to *Schrodinger*’s *Simulation Interactions Diagram* to build plots and related metrics for analysis, such as Root Mean Square Fluctuation (RMSF), Root Mean Square Deviation (RMSD), among others. Additionally, RMSD boxplots as well as merged RMSF graphs were generated using our in-house script available at the https://github.com/guimsilvaa/desmotools repository.

The *MMGBSA* ΔG free energy binding affinities values between protein-ligand complexes were calculated for each MD trajectory-frame by *Schrodinger*’s *thermal_mmgbsa.py* script. These were calculated using the OPLS4 force field and the default *Prime* tool. Statistical parameters such as average and corresponding standard deviation values for ΔG and associated components were also calculated using available scripts also at our *desmotools* repository. For the structural comparison of SLC25A45 and SLC25A48, SLC25A48 structure was retrieved from AlphaFoldDB under the code: AF-Q6ZT89-F1-model_v4 and prepared using same procedure as described above. Sequence alignment was carried out using Multiple sequence viewer and Muscle algorithm from Maestro. Structure alignment was performed using Matchmaker from ChimeraX 1.9. All figures were generated using *Pymol* 3.0.2.

### Mitochondrial respiration (*JO_2_*)

Tissues or cells were homogenized in MHSE buffer (70 mM sucrose, 210 mM mannitol, 5 mM HEPES, 1 mM EGTA, pH 7.2) plus 0.5 % (w/v) fatty-acid free BSA using a motorized Teflon glass homogenizer. The homogenate was centrifuged at 600 x g for 10 min at 4°C. Following the removal of fat and lipid by aspiration, the supernatant was centrifuged at 9,000 x g for 10 min at 4C to obtain a mitochondrial pellet. After resuspending the pellet in MHSE + BSA, the 9,000 x g centrifugation step was repeated, and the final pellet was resuspended in a minimal volume of MHSE. The protein concentration was determined using the bicinchoninic assay (Pierce). Respiration was measured using an Oroboros O2k respirometer (Oroboros Instruments, Austria) with mitochondria suspended in Buffer Z (1 mM EGTA, 5 mM MgCl_2_, 105 mM K-MES, 30 mM KCl, 10 mM KH_2_PO_4_, 5 mg/mL fatty acid-free BSA, pH 7.1). Complex I-driven respiration was initiated with 10 mM glutamate, 2 mM malate and 5 mM pyruvate, while complex II-driven respiration was fueled with 10 mM succinate. Beta-oxidation was measured via the addition of 25 µM palmitoyl-CoA, 2 mM L-carnitine and 0.5 mM malate. State 3 respiration was stimulated with 4 mM ADP. To determine the contribution of UCP1 to respiration in brown adipose tissue, 4 mM GDP was added to the chamber.

### Mouse physiology

Lean and fat mass were measured via the 3-in-1 Echo MRI Composition Analyzer (Echo Medical Systems) prior to and after a 7-day treatment with semaglutide (30 nmol/kg). For intraperitoneal glucose tolerance tests, mice were fasted for 6 h prior to an injection (1.5 g/kg) of glucose. For insulin tolerance tests, mice were fasted for 3 h and insulin (0.5 U/kg body weight) was injected. Blood samples for glucose measurements were taken at 15, 30, 60, 90, and, 120 min post-injection using blood glucose test strips (Freestyle Lite). For acute cold tolerance tests, eight-week-old *Slc25a45*^CMV^KO or *Slc25a45*^Alb^KO mice and their respective littermate controls were housed at thermoneutrality (30°C) for 1 week prior to cold exposure at 6°C for 5 hours in the absence of food. To test if carnitine supplementation restores cold sensitivity in SLC25A45 Ko mice, a cohort of *Slc25a45*^CMV^KO mice was given L-carnitine (100 mg/kg) for five days prior to being placed in the cold. Hourly core temperatures were taken using a TH-5 rectal thermometer (Physitemp). For other cold studies, such as serum metabolomics and lipidomics studies, mice were group-housed with their littermate controls and provided with food and enrichment to prevent mortality due to hypothermia. Mice housed at room temperature were gradually acclimated to cold temperatures at 6°C for 72 hours.

We used the following methods to quantify metabolites and hormone levels. L-carnitine in mouse tissues and serum was measured as per the manufacturer’s protocol (Colorometric assay; Abcam, ab83392). Serum insulin concentrations were measured after a 6 h fast according to the manufacturer’s recommendations (Millipore; EZRMI-13K). For kidney damage markers, serum creatinine and blood urea nitrogen were probed per the manufacturer’s protocols (MyBioSource, MBS763433 and MBS2611085, respectively). Cystatin C was measured following the directions in the kit (proteintech; KE10066). Tissue malondialdehyde was measured to indicate lipid peroxidation in the kidney using a commercially available kit (Colorometric assay; Abcam, ab118970). Free glycerol in circulation was measured with 10 μL of serum in duplicate per the manufacturer’s protocol (Sigma-Aldrich, F6428).

### Indirect calorimetry

Whole-body energy expenditure (VO_2_, VCO_2_), respiratory exchange ratio, food intake, and locomotor activity (beam break counts) of *Slc25a45* KO mice and littermate control mice were monitored using the Promethion Metabolic Cage System (Sable Systems) by the BIDMC Energy Balance Core. Mice had unrestricted access to food and water throughout the study. The obtained data were analyzed by CalR (https://bankslab.shinyapps.io/prod_ver/).

### Plasma lipidomics

Samples were extracted in butanol/methanol (1:1) with 5 mM ammonium formate. Chromatographic separation was conducted using a Vanquish UHPLC (Thermo Scientific, Waltham, MA, USA) and Acquity CSH C18 column (1.7 µm, 2.1 mm × 100 mm) with guard column (1.7 µm, 2.1 mm × 5 mm) (Waters, Milford, MA). Mobile phase A was water: acetonitrile 40:60, and mobile phase B isopropanol: acetonitrile 90:10, both mobile phases containing 10 mM ammonium formate and 0.1% formic acid. The elution gradient was 20% B from 0 to 3 min, 55% B at 7 min, 65% B at 15 min, 70% B at 21 min, 88% B at 23 min, 100% B at 24 min held until 26 min, and 20% B at 28 min and held until 30 min. The flow rate was 0.35 mL/min. The autosampler was at 4°C. The injection volume was 5 µL. Needle wash was performed between samples using dichloromethane: isopropanol: acetonitrile at 1: 1: 1. The mass spectrometry was Exploris 480 orbitrap (Thermo Scientific, Waltham, MA, USA). The ion source was H-ESI. Spray voltage was 3500 V for positive ions, and 2500 V for negative ions. Sheath gas was set at 50 arbitrary unit (Arb), auxiliary gas 15 Arb, sweep gas 1 Arb, ion transfer tube at 325 °C, and vaporizer 350 °C. The scan mode was data dependent (dd)-MS2, covering 150-1600 m/z in both positive and negative polarities. Precursor ion scan had resolution 60,000. For product ion scan, the resolution was 15,000, isolation width 1.0 m/z, and collision energy 25%. Thermo Scientific LipidSearch software version 5.0 was used for lipid identification and quantitation. First, the product search mode was used to identify lipids based on the exact mass of the precursor ions and the MS2 mass spectra of the product ion scan. The precursor and product tolerance was 10 ppm. The absolute intensity threshold of precursor ions and the relative intensity threshold of product ions were set to 30000 and 1%, respectively. Next, the search results from all samples were aligned within a retention time tolerance of 0.25 min. The annotated lipids were then filtered to reduce false positives by only including the lipids with a total grade of A and B.

### Tissue histology and image acquisition

Kidney and inguinal white adipose tissues were fixed in 4% paraformaldehyde overnight at 4°C, followed by dehydration in 70% ethanol. After the dehydration procedure, tissues were embedded in paraffin and cut into sections at a thickness of 5 μm. The sections were processed for H&E staining according to the standard protocol at the BIDMC pathology core. Images were acquired using a Revolve microscope (ECHO Laboratories). A scalebar (200 μm) was added to each image. Adipocyte area was measured from three biological replicates per group using the Adiposoft plugin for Fiji.

### Transmission electron microscopy

Tissues were immersion fixed in 2% glutaraldehyde (Electron Microscopy Sciences, Hatfield, PA) and 2.5% paraformaldehyde (Electron Microscopy Sciences, Hatfield, PA) in 0.1M sodium cacodylate (Sigma-Aldrich, Burlington, MA) pH 7.4 for at least 1 h at 23°C and then at 4°C overnight. Tissues were washed with 0.1M sodium cacodylate and then post-fixed for 1 h at 4°C in 1% osmium tetroxide (Electron Microscopy Sciences) in 0.1M sodium cacodylate. Cells were washed in DI water and incubated in 2% aqueous uranyl acetate (Electron Microscopy Sciences) overnight at 4°C. The next day, tissues were washed with DI water and then dehydrated at 4°C in a graded ethanol series. Tissues were then brought to 23°C and dehydrated with 100% ethanol (Sigma-Aldrich) followed by propylene oxide (Electron Microscopy Sciences). Infiltration with LX112 resin (Ladd Research Industries, Williston, VT) was followed by embedding in flat-bottom Beem capsules (Electron Microscopy Sciences). The resulting blocks were sectioned using a Leica Ultracut E ultramicrotome (Leica Microsystems, Wetzlar, Germany) and sections placed on formvar and carbon-coated grids (Electron Microscopy Sciences). The sections were contrast-stained with 2% uranyl acetate followed by lead citrate (Sigma-Aldrich), and imaged in a JEOL 1400 transmission electron microscope (JEOL, Peabody, MA) equipped with a Gatan Orius SC1000 digital CCD camera (Gatan, Pleasanton, CA).

### Single-nuclei RNA-seq

Data from GSE164274 were analyzed and visualized using R (v4.0.4) and Seurat (v4.0.1) following standard workflows.

### RNA-seq

Total RNA was purified using TRIZOL (ThermoFisher). RNA samples were quantified using a Qubit 2.0 Fluorometer (Life Technologies, Carlsbad, CA, USA), and RNA integrity was checked with the 4200 TapeStation (Agilent Technologies, Palo Alto, CA, USA). ERCC RNA Spike-In Mix (ThermoFisher) was added to normalized total RNA prior to library preparation following the manufacturer’s protocol. Next, we performed rRNA depletion using the QIAGEN FastSelect rRNA HMR Kit (Qiagen). RNA sequencing libraries were constructed with the NEBNext Ultra II RNA Library Preparation Kit for Illumina according to the manufacturer’s instructions. The sequencing libraries were multiplexed and clustered on the flowcell. After clustering, the flowcell was loaded on the Illumina NovaSeq instrument. The samples were sequenced using a 2x150 Paired-End (PE) configuration. After demultiplexing, sequence data was checked for overall quality and yield. Then, raw sequence reads were trimmed to remove possible adapter sequences and nucleotides with poor quality using Trimmomatic v.0.36. The reads were then mapped to the reference genome (mm10) using the STAR aligner v.2.5.2b. Unique gene hit counts were calculated by using feature Counts from the Subread package v.1.5.2. Only unique reads that fell within exon regions were counted. Downstream analysis was performed on R or python. PCA is visualized with python. Using edgeR, a comparison of gene expression between the groups of samples was performed. Genes with a false-discovery rate of less than 0.05 were called as differentially expressed genes. A heatmap was drawn using pheatmap on R. Pathway enrichment was evaluated using the Mann–Whitney U test on gene-level log₂ fold changes in Python, followed by Benjamini–Hochberg correction.

### Quantitative RT-PCR

Trizol (Invitrogen) was used to isolate total RNA from cells or tissue according to manufacturer instructions. The iScript cDNA synthesis kit (Biorad) was used to reverse transcribe RNA. PCR reactions were performed on the QuantStudio 6 Flex Real Time PCR System (Applied Biosystems) using SYBRgreen (Bio-Rad). Assays were performed in duplicate, and all results were normalized to 18S ribosomal RNA (human) or 36B4 (mouse). Values are relative to the mean of the control group. Primers used are listed in **Table S2**.

### Statistics and reproducibility

Data were expressed as mean ± s.e.m and statistical analysis was performed using GraphPad Prism 8 (GraphPad Software, Inc., La Jolla, Ca) unless otherwise mentioned. P < 0.05 was considered statistically significant throughout the study. When two-group comparisons were performed, a two-sample unpaired Student’s t-test was used. For multiple group comparisons, we used a one-way ANOVA followed by Tukey’s post hoc HSD test. Two-way ANOVA with Šídák’s multiple comparisons test was used for cold tolerance tests and cellular D_9_-TML uptake time courses. Kidney and adipose histology were performed blinded, and representative images are shown from 3 biological samples per group.

## ACKNOWLEDGMENTS

We are grateful to Dr. David J Pagliarini for sharing the bacterial expression system for the transporter assays, and Tadashi Yamamuro in the Kajimura lab for their technical support.

## Author Contributions

Conceptualization: CA, SK. Methodology: CA, HN, BY, GMS, ML, DW, ASB, LS. Investigation: CA, HN, BY, GMS, MF, DK, DW, MGP, JS, RW, JY, ARPV. Visualization: CA, GMS, LS. Funding acquisition: ASB, SH, SK. Supervision: SH, LS, SK. Writing-original draft: CA, SK. Writing-reviewing & editing: all authors.

## Data and materials availability

The raw sequencing data generated in this study have been deposited in the NCBI Sequence Read Archive (SRA) under the BioProject accession number PRJNA1345061.All LC-MS raw data for metabolomics have been uploaded to the Metabolomics Workbench (https://www.metabolomicsworkbench.org), with the project ID ST003920 (http://dx.doi.org/10.21228/M8JZ5D). All other data are available in the manuscript or supplementary materials. All the materials and mice are available after completion of MTA.

---

**Fig. S1.**
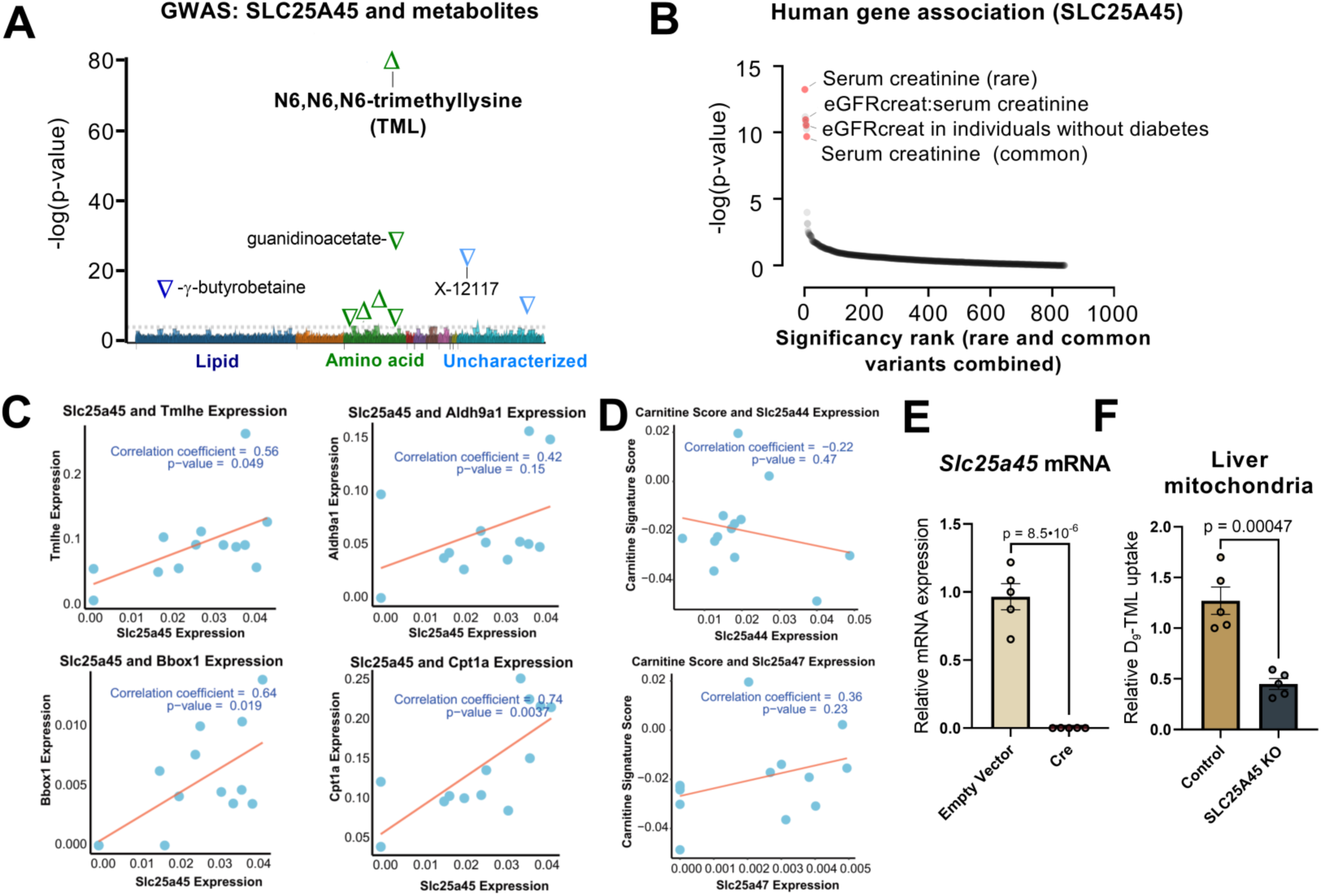
SLC25A45 is required for L-carnitine synthesis from trimethyllysine (TML). **(A)** GWAS-METSIM of 6,136 participants for circulating metabolites associated with *SLC25A45.* TML was the top hit with a p-value of *P* = 1 × 1.9^-80^. **(B)** Genetic association of *SLC25A45* for human diseases and traits available in the common metabolic disease knowledge portal (https://hugeamp.org). **(C)** Correlation between *Slc25a45* expression and enzymes related to carnitine biosynthesis and fatty acid oxidation. Statistics: Pearson correlation test. **(D)** Correlation between mitochondrial solute carrier (*Slc25a44* and *Slc25a47*) expression and carnitine signature score. **(E)** Relative mRNA expression of *Slc25a45* in inguinal white adipose-derived stromal vascular fraction. n = 5 per group. Statistic: unpaired t-test. **(F)** TML uptake in isolated mitochondria of control and SLC25A45 KO mice. n = 5 per group. Statistic: unpaired t-test.

**Fig. S2.**
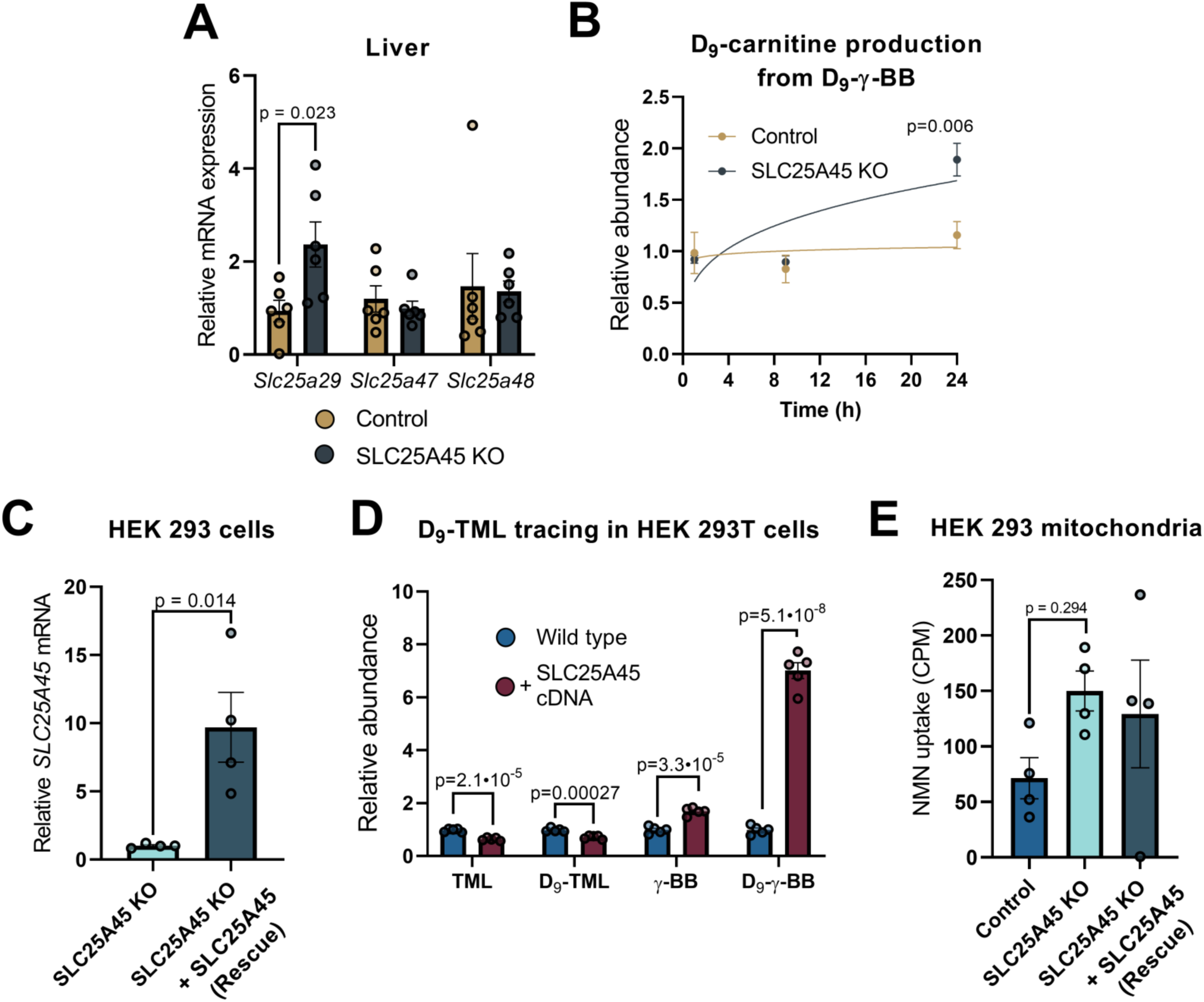
SLC25A45 facilitates the synthesis of L-carnitine from trimethyllysine. **(A)** Relative mRNA expression of mitochondrial solute carriers in liver tissue from control and SLC25A45 KO mice. n = 6 per group. Statistic: unpaired t-test. **(B)** D_9_-labeled γ-butyrobetaine (γ-BB) tracing at 10 µM in hepatocytes derived from control and SLC25A45 KO mice. n = 3 per group. Statistic: Two-way ANOVA with Šídák’s multiple comparisons test. **(C)** Relative mRNA expression of *SLC25A45* in SLC25A45 KO HEK 293T cells and SLC25A45 KO cells overexpressing cDNA for SLC25A45 (Rescue). n = 4 per group. Statistic: unpaired t-test. **(D)** D_9_-labeled TML tracing in WT HEK293 and WT cells expressing ectopic SLC25A45. n = 5 per group. Statistic: unpaired t-test. **(E)** Mitochondrial uptake of tritium-labelled nicotinamide mononucleotide in control, SLC25A45 KO HEK 293T cells, and SLC25A45 KO cells overexpressing SLC25A45 (Rescue). n = 4 per group. Statistic: One-way ANOVA with Tukey’s post-hoc HSD test.

**Fig. S3.**
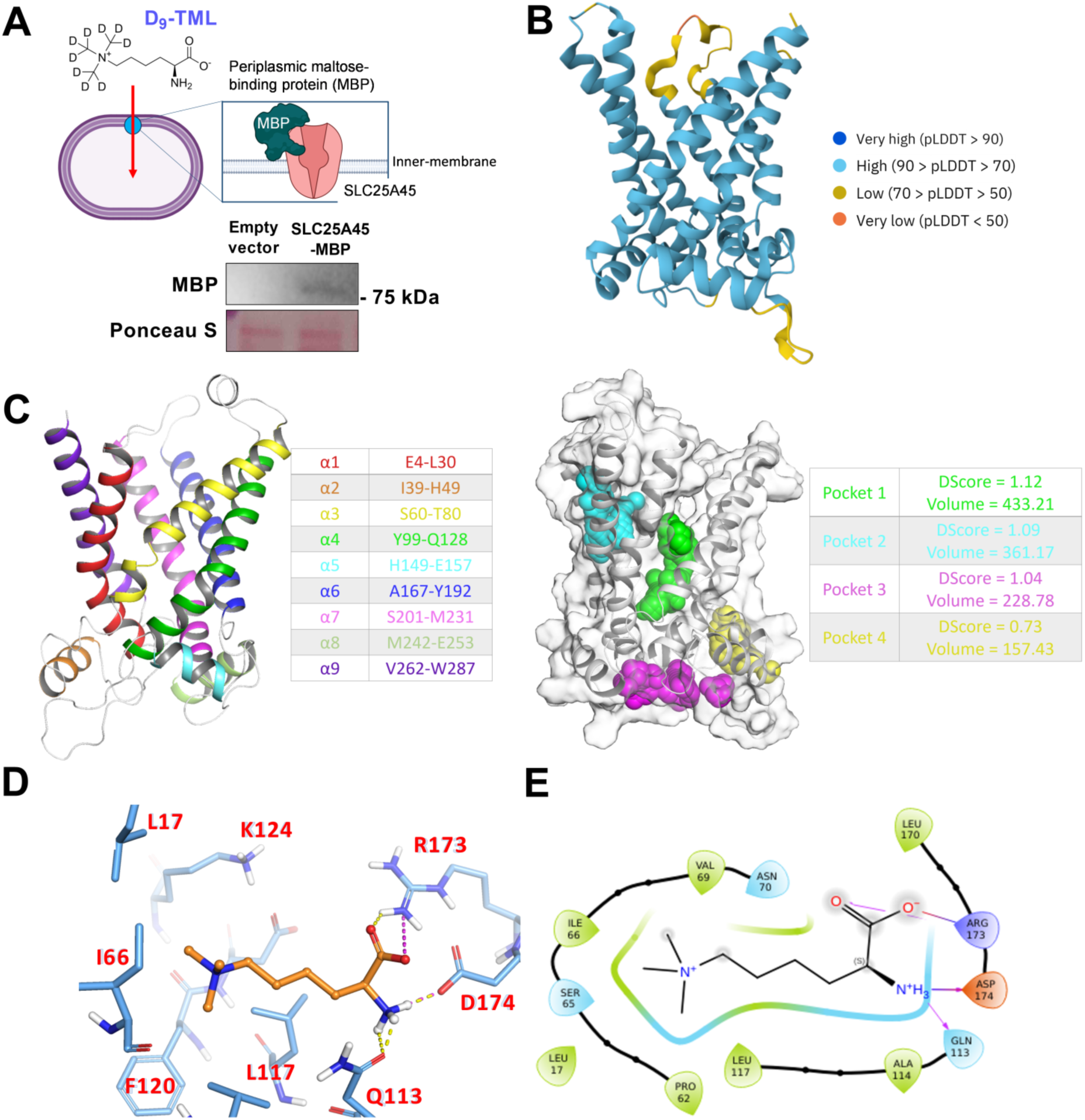
SLC25A45 transports TML in the mitochondria. **(A)** Schematic of MBP-SLC25A45-expressing *E.coli* and representative immunoblotting of MBP-SLC25A45. **(B)** AlphaFold structure (AF-Q3KQZ1-F1-model_v4) for SLC25A45. **(C)** Cartoon/ribbon representation of SLC25A45 prepared structure showing its nine (six in vertical – α1, α3, α4, α6, α7, and α9) transmembrane alpha helices and corresponding range of amino acids, in respective coloring scheme. (b) Surface and cartoon (in white) representation of SLC25A45 prepared structure and detected cavities (pockets) by *Sitemap*. **(D)** A 3D view of the docking pose was obtained for TML (orange) key interactions (dashed lines in yellow and magenta correspond to the hydrogen bond and salt bridge, respectively), which was established with SLC25A45 key residues (blue). **(E)** 2D interaction diagram for TML-SLC25A45, where magenta arrows and solid lines represent hydrogen bonds and salt bridges, respectively.

**Fig. S4.**
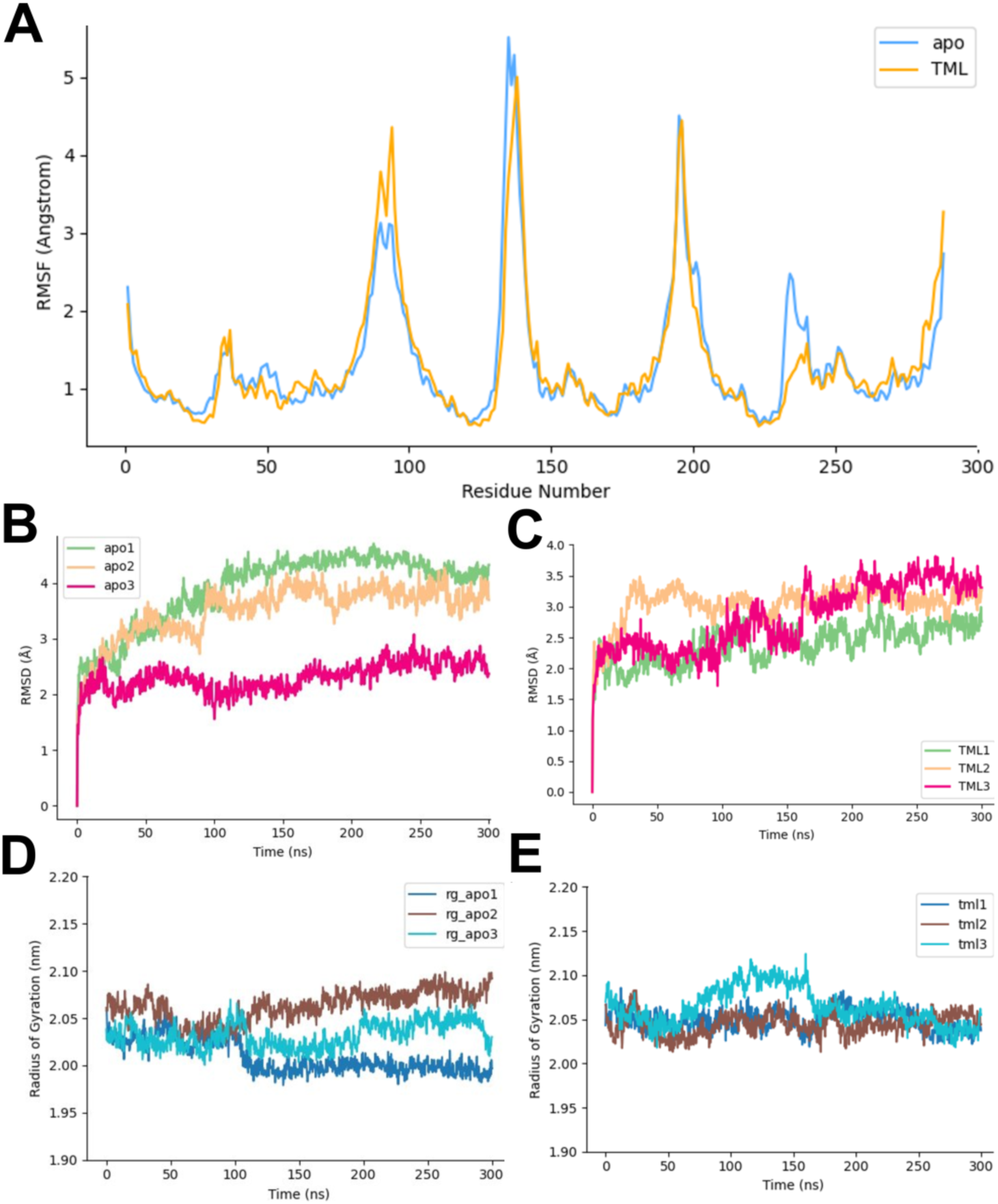
MD simulations for SLC25A45 *apo* and complexed with TML. **(A)** Average root mean square fluctuation values (in angstroms) for SLC25A45 amino acid residues, with (yellow) and without (apo, blue) TML. **(B)** Root mean square deviation values (in angstroms) obtained for each 300 ns simulation replicate of the SLC25A45 protein backbone in its apo form (without ligand). **(C)** Root mean square deviation values (in angstroms) obtained for each 300 ns simulation replicate of the SLC25A45 protein backbone when bound to TML. **(D)** Radius of gyration values (in nanometers) obtained for each 300 ns simulation replicate of the SLC25A45 in its apo form (without ligand). **(E)** Radius of gyration values (in nanometers) obtained for each 300 ns simulation replicate of the SLC25A45 when bound to TML.

**Fig. S5.**
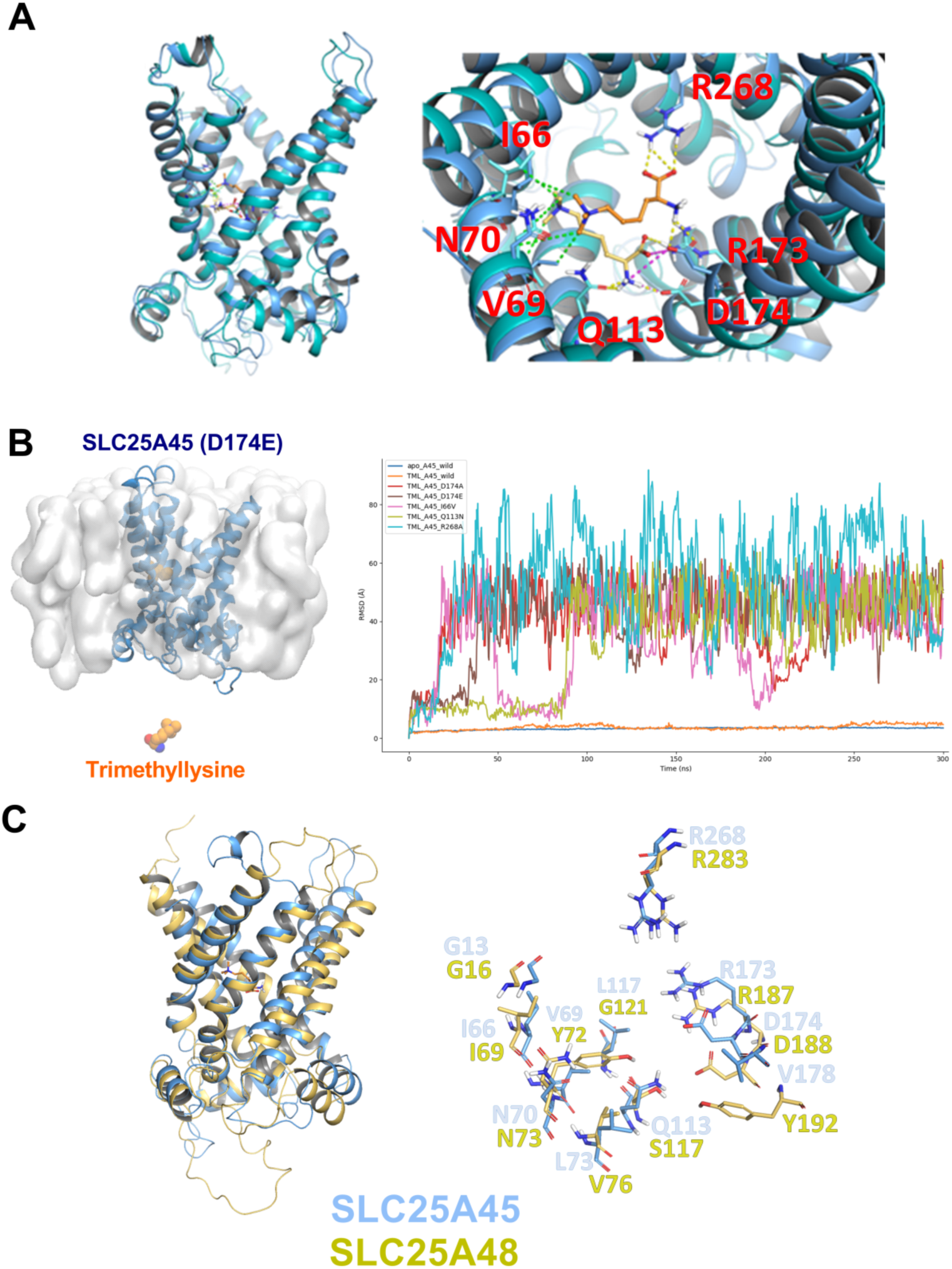
Structural insights into the substrate specificity for SLC25A45. **(A)** Structural views of the SLC25A45-TML complex before and after MD simulations. Left: Side view of the A45-TML complex. The “before” structure is shown in teal (protein) and yellow (TML), while the “after” structure is shown in dark blue (protein) and orange (TML). Right: Top-zoom view of the A45-TML complex highlighting the hydrophobic interactions (green dashed lines) between TML and residues I66, N70, and V69 before/after MD. Additional interactions include hydrogen bonds and salt bridges represented by dashed lines in yellow and magenta, respectively, and key interacting residues are labeled for clarity. **(B)** Left: Side view of SLC25A45 (D174E) at the inner mitochondrial membrane and TML escaping the inner cavity. Right: Average RMSD (protein backbone) against time (ns) plot obtained by MD simulations for SLC25A45 *apo* (blue), complexed with TML (orange), and various mutants demonstrating less stable interactions between the protein and ligand. **(C)** Left: Structural alignment of SLC25A45 (blue) and SLC25A48 (yellow), highlighting the overall similarity in their 3D conformations. Right: Close-up view of key residues surrounding the hydrophobic N-trimethyl moiety in the SLC25A45-TML complex. Pairs of conserved (e.g., I66/I69, N70/N73) and non-conserved (e.g., V69/Y72, L73/V76, L117/G121) residues are labeled.

**Fig. S6.**
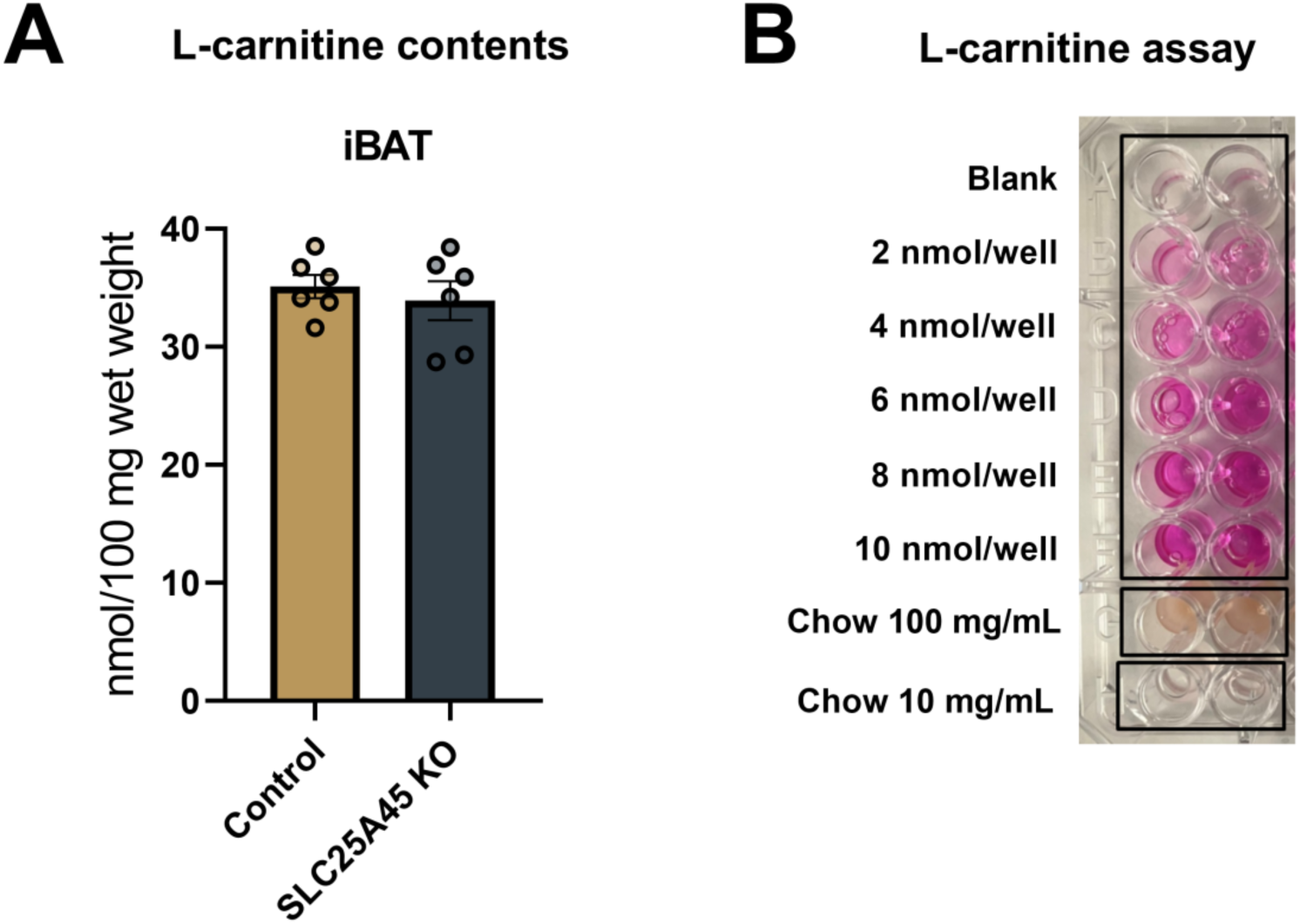
L-carnitine contents of tissue and diet. **(A)** Carnitine contents in the interscapular brown adipose tissue (iBAT) from control and SLC25A45 KO n=6 per group. Statistic: unpaired t-test. Bars represent mean and error, as shown in s.e.m. **(B)** Representative image of spectrophotometric L-carnitine assay demonstrating a standard curve and murine chow diet dissolved in assay buffer. The dietary carnitine content at 570 nm was below the detection limit.

**Fig. S7.**
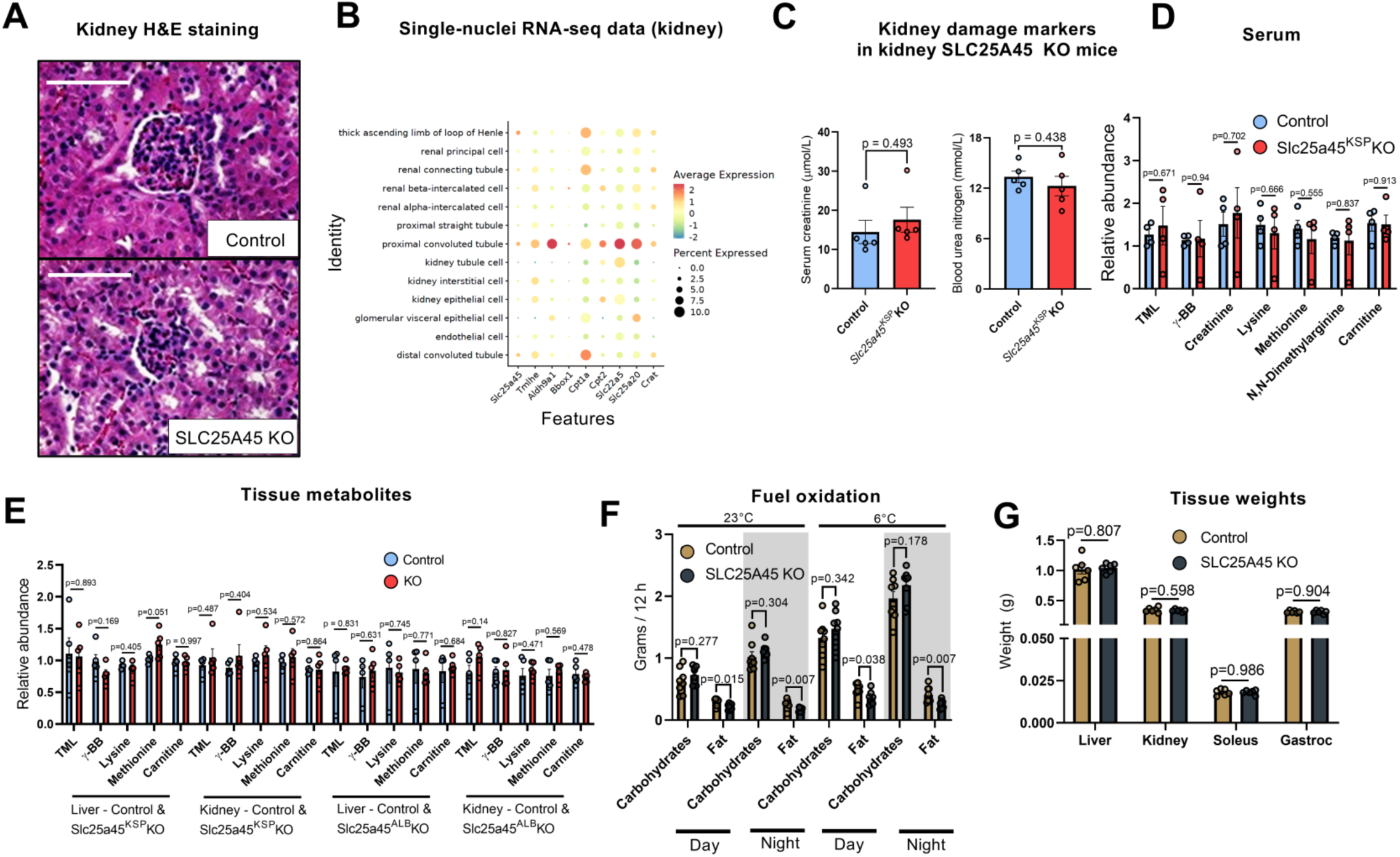
Characterization of whole-body and tissue-specific SLC25A45 KO mice. **(A)** Representative H&E staining of kidney sections from Control and whole-body SLC25A45 KO mice. Scale bar = 200 µm. **(B)** Single-nuclei RNA-seq data of indicated genes involved in the *de novo* biosynthesis of carnitine and the carnitine shuttle in the kidney. **(C)** Serum levels of kidney damage markers creatinine and blood urea nitrogen from control and kidney-specific SLC25A45 KO mice. n=5 per group. Statistic: unpaired t-test. **(D)** Serum levels of TML, carnitine, and related metabolites in control and kidney-specific SLC25A45 KO mice. n = 5 per group. **(E)** Tissue levels of TML, carnitine, and related metabolites in control, liver and kidney specific KO mice. n = 5 per group. **(F)** Estimated carbohydrate (*CHO*_ox_ = (4.585x*V*CO_2_) – (3.226x*V*O_2_)) and fat oxidation (*FAT*_ox_ = (1.695x*V*O_2_) – (1.701x*V*CO_2_) derived from metabolic caging data in control and SLC25A45 KO mice. n = 8 per group. Statistic: unpaired t-test. **(G)** Tissue weights from 30-week-old control and SLC25A45 KO mice. n = 6 per group.

**Fig. S8.**
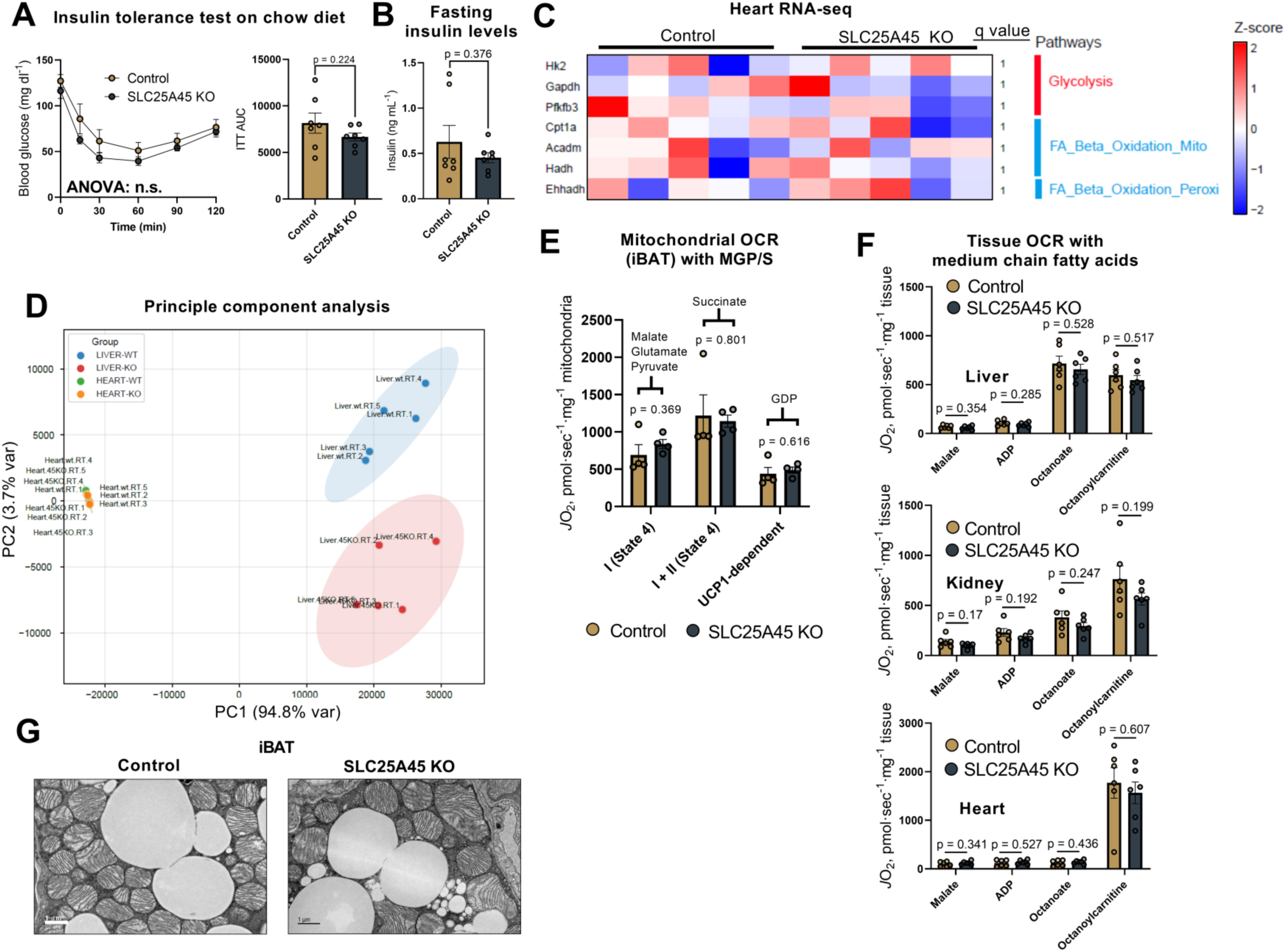
Compensatory metabolic pathways in whole-body SLC25A45 KO mice. **(A)** Insulin tolerance test of male mice after a six-hour fast. n=7. AUC statistic: unpaired t-tests. **(B)** Fasting insulin levels after a six-hour fast. n=7. Statistic: unpaired t-test. **(C)** Relative mRNA levels of indicated genes in the heart at 23℃. Data represented as a Z-score heatmap for each gene in each sample, representing the quantified value. Pathways for each gene are listed on the right. n = 5. Statistic: the Benjamini-Hochberg procedure in R. **(D)** Principal component (PC) analysis of heart and liver based on their transcriptomic profiles. **(E)** Mitochondrial respiration (*J*O_2_) in the iBAT in the presence of glutamate, malate, and pyruvate (Complex I) and Complex II-driven respiration with succinate. UCP1-dependent respiration was determined with GDP. n=4. Statistic: unpaired t-test. **(F)** Mitochondrial respiration in the liver, kidney, and heart in response to medium-chain fatty acid (C8, octanoate) or acylcarnitine (octanoylcarnitine) in the presence of malate, ADP, octanoate, and octanoylcarnitine. n = 6. **(G)** Representative transmission electron microscopy images showing mitochondrial structure from iBAT. Scale bar = 1 µm.

**Fig. S9.**
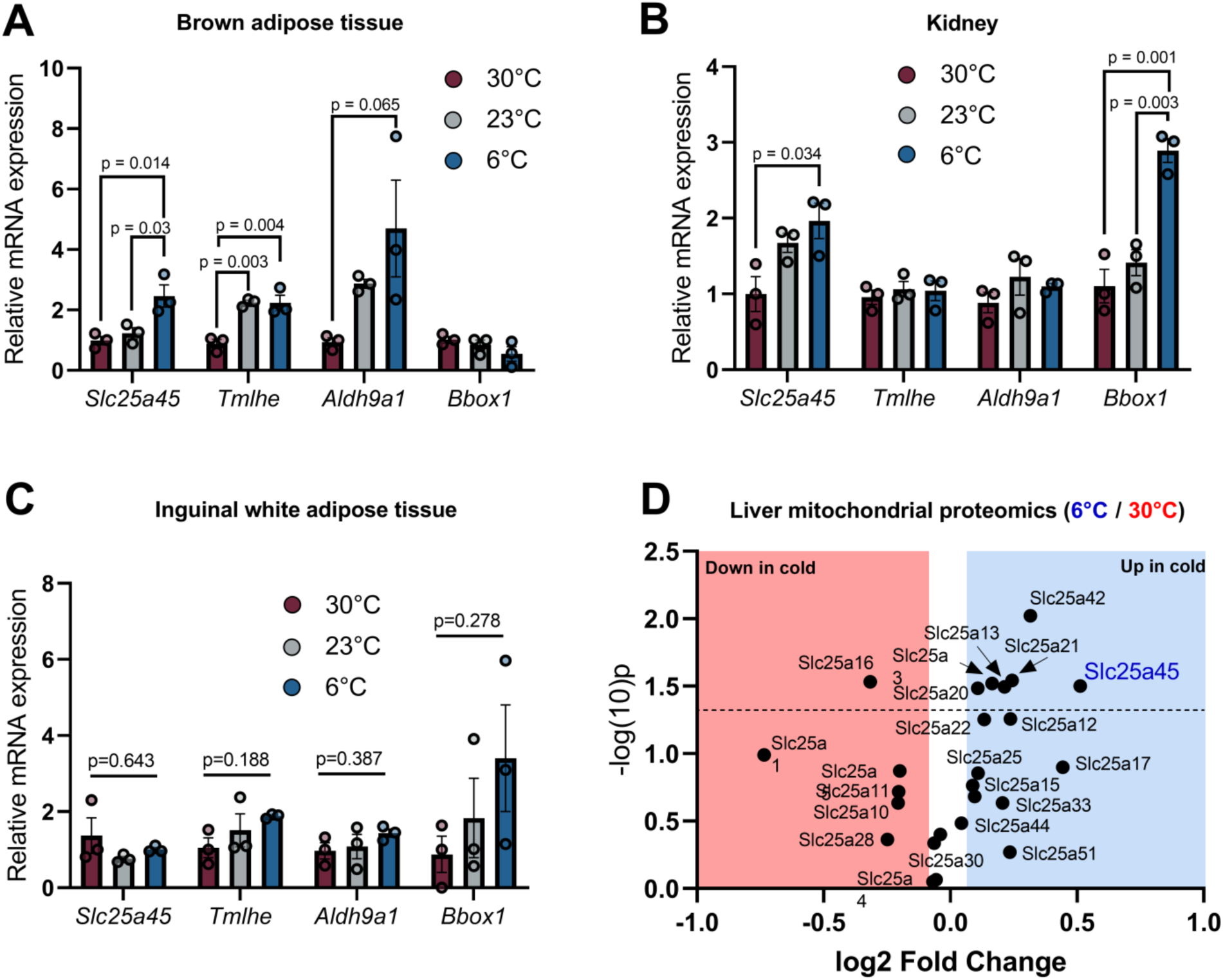
SLC25A45 expression is induced by cold exposure. **(A)** Relative mRNA expression of indicated carnitine synthesis genes in the brown adipose tissue of wild-type male C57BL/6J mice acclimated to thermoneutrality, room temperature, and 6°C for 72 h. n = 3 per group. Statistic: One-way ANOVA with Tukey’s post hoc HSD test. **(B)** Relative mRNA expression of indicated carnitine synthesis genes in the kidney of wild-type male C57BL/6J mice. n = 3 per group. Statistic: One-way ANOVA with Tukey’s post hoc HSD test. **(C)** Relative mRNA expression of indicated carnitine synthesis genes in the inguinal white adipose tissue of wild-type male C57BL/6J. *n* = 3 per group. Statistic: One-way ANOVA with Tukey’s post hoc HSD test. **(D)** Volcano plot of liver mitochondrial proteomics detected changes in the expression of mitochondrial solute carriers in mice acclimated to 6 or 30°C. The data are from Liu, Q et al^34^. *n* = 3 per group.

**Fig. S10.**
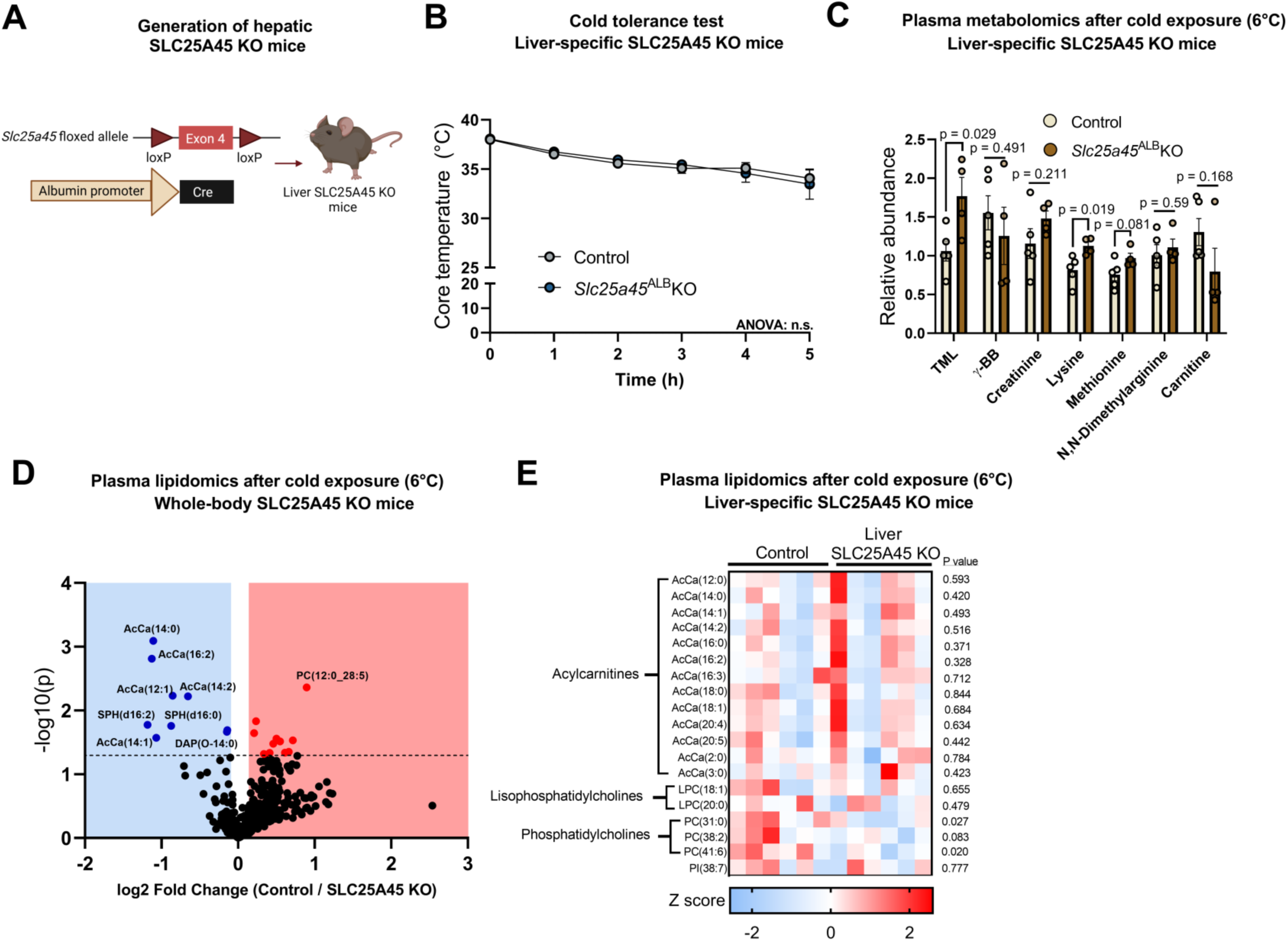
Serum metabolites and lipid profile of liver-specific SLC25A45 KO mice. **(A)** Schematic demonstrating the development of liver-SLC25A45 KO mice using *Alb*-Cre. **(B)** Cold tolerance test of liver-specific SLC25A45 KO mice and littermate control mice. n=7 per group. Statistic: two-way ANOVA with Šídák’s multiple comparisons test. **(C)** Serum levels of TML, carnitine, and related metabolites in control and liver-specific SLC25A45 KO mice. n = 5 per group. Statistic: unpaired t-test. **(D)** Volcano plot of serum lipidomics from whole-body SLC25A45 KO mice and littermate controls, group-housed at 6°C for 72 h. **(E)** Serum lipidomics analysis of liver-specific SLC25A45 KO mice and littermate controls following cold exposure for 72h. n=6 per group. Statistic: unpaired t-test. Data represented as Z score.

**Fig. S11.**
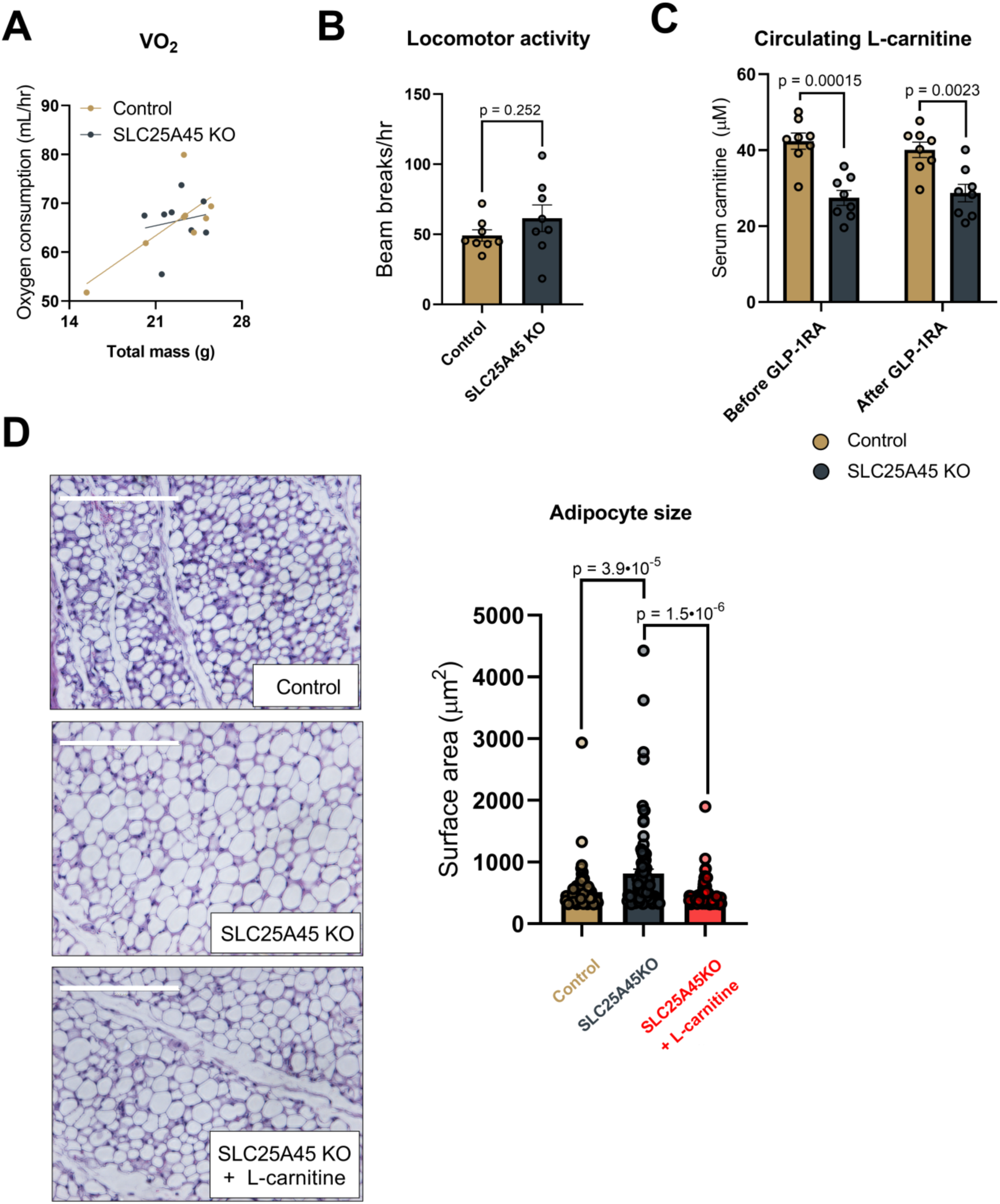
SLC25A45 is required for GLP-1RA-induced adipose tissue loss. **(A)** Daily whole-body oxygen consumption rate (VO_2_, ml hr^-1^) from Fig. 6A by CaIR-ANCOVA. n=8. **(B)** Locomotor activity from Fig. 6A. n=8. Statistic: unpaired t-test. **(C)** Serum carnitine levels before and one day after an injection of semaglutide at 30 nmol kg^-1^. n = 8. Statistic: unpaired t-test. **(D)** Representative H&E images (left). Mice were treated with semaglutide as shown in Fig. 6H-M. Scale bar = 200 µm. Right: Adipocyte size was measured using the Adiposoft plugin for Fiji. n=89 per group. Statistic: One-way ANOVA with Tukey’s post hoc HSD test.

**Table S1.** MMGBSA ΔG binding affinity estimation for three replicates of MD simulations performed for the complex of TML bound at the SLC25A45 binding site.

| | MMGBSA $\Delta G$ (kcal/mol) | | | |
| --- | --- | --- | --- | --- |
|  | replicate1 | replicate2 | replicate3 | average |
| **SLC25A45**<br>**+**<br>**TML** | $-14.35 \pm 3.16$ | $-17.39 \pm 3.50$ | $-19.57 \pm 4.60$ | **$-17.10 \pm 3.75$** |

**Table S2.**
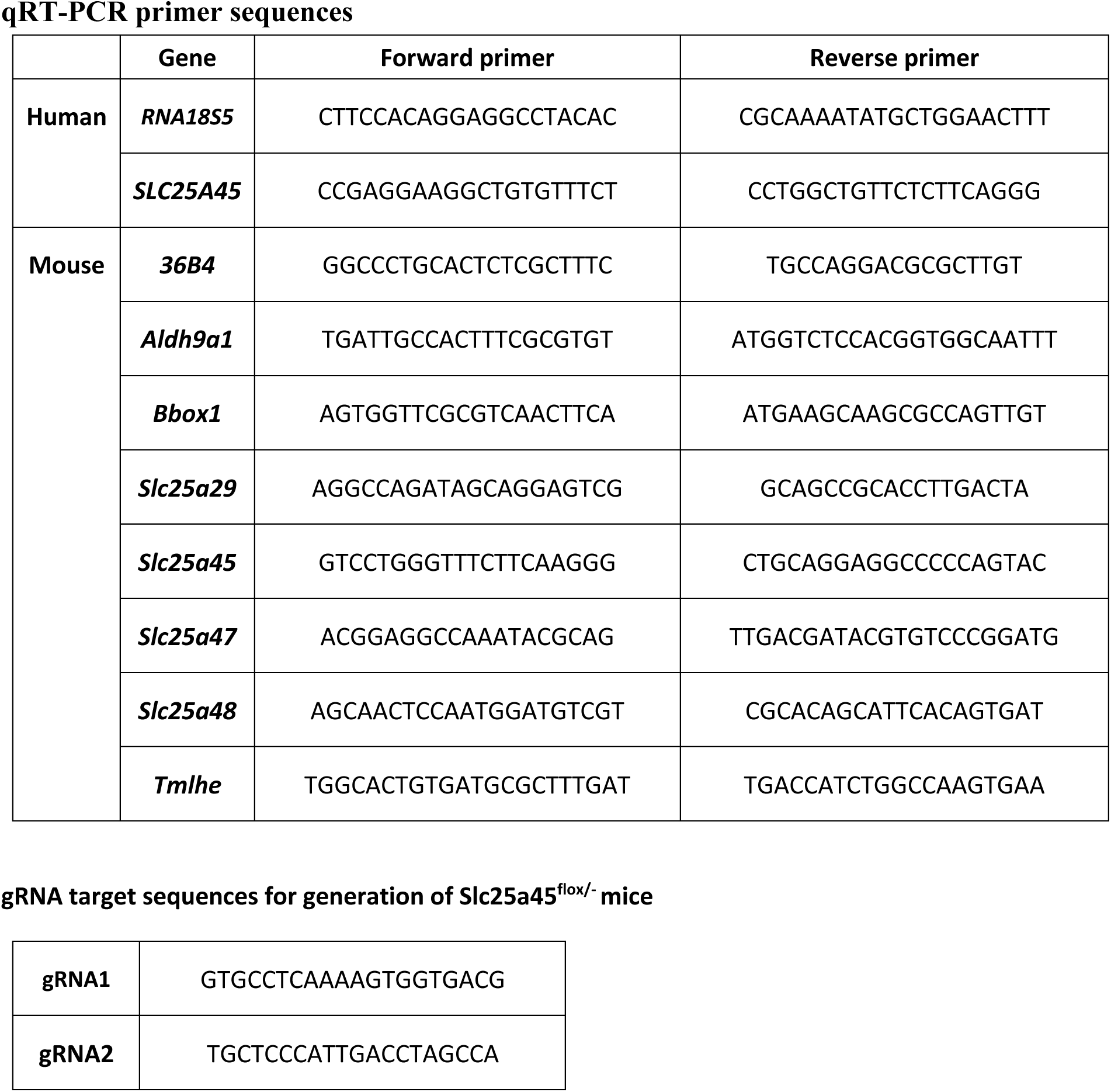
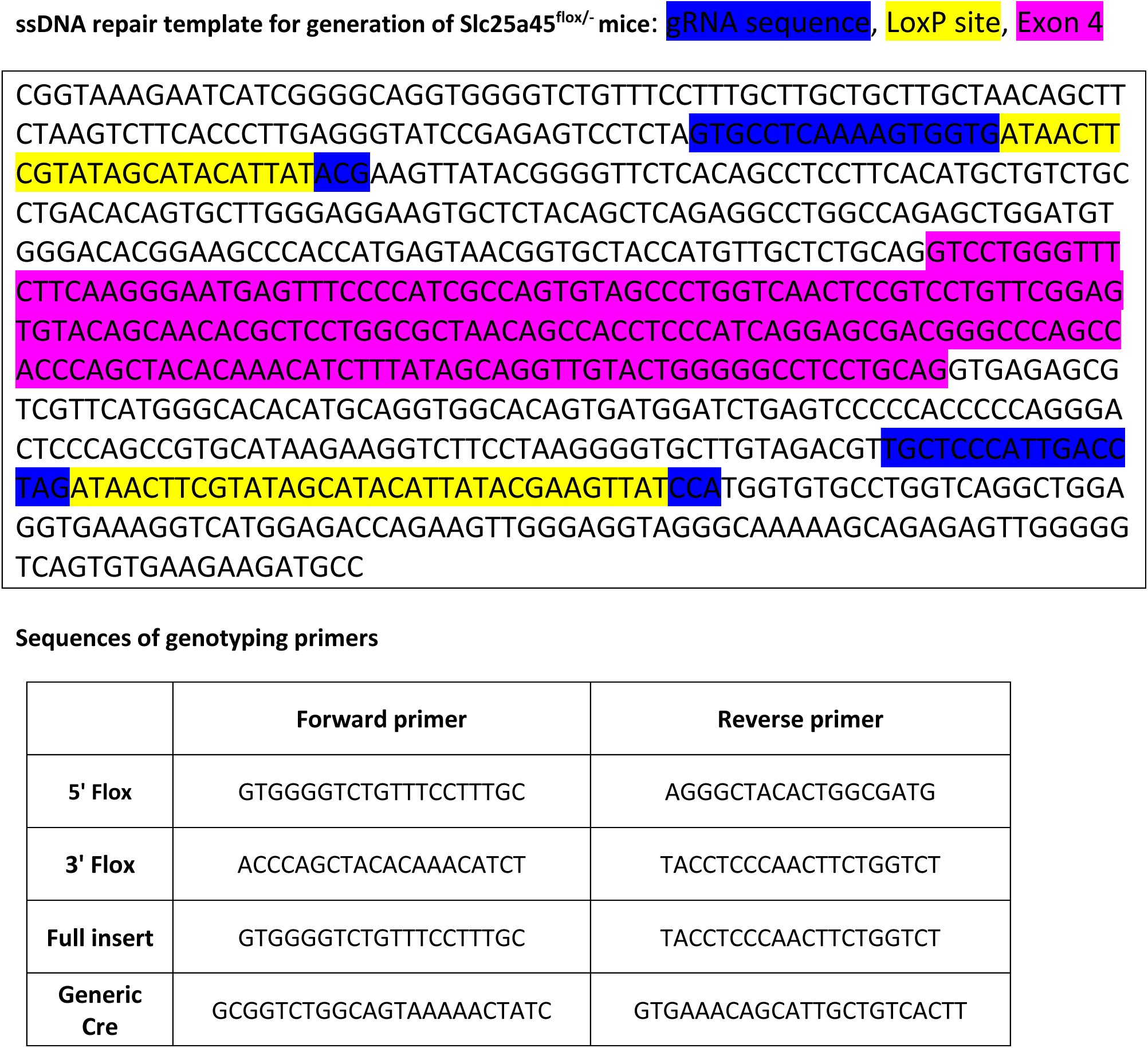
qRT-PCR primer sequences.

## REFERENCES

1. B. H. Goodpaster, L. M. Sparks, Metabolic Flexibility in Health and Disease. Cell metabolism 25, 1027–1036 (2017).

2. R. L. Smith, M. R. Soeters, R. C. I. Wust, R. H. Houtkooper, Metabolic Flexibility as an Adaptation to Energy Resources and Requirements in Health and Disease. Endocrine reviews 39, 489–517 (2018).

3. M. R. Bornstein et al., Comprehensive quantification of metabolic flux during acute cold stress in mice. Cell metabolism 35, 2077–2092 e2076 (2023).

4. T. Yoneshiro et al., BCAA catabolism in brown fat controls energy homeostasis through SLC25A44. Nature 572, 614–619 (2019).

5. A. R. P. Verkerke et al., BCAA-nitrogen flux in brown fat controls metabolic health independent of thermogenesis. Cell 187, 2359–2374 e2318 (2024).

6. G. D. Lopaschuk, Q. G. Karwi, R. Tian, A. R. Wende, E. D. Abel, Cardiac Energy Metabolism in Heart Failure. Circulation research 128, 1487–1513 (2021).

7. F. M. Vaz, R. J. Wanders, Carnitine biosynthesis in mammals. The Biochemical journal 361, 417–429 (2002).

8. A. C. Ross, B. Caballero, R. J. Cousins, K. L. Tucker, T. R. Ziegler, Modern nutrition in health and disease: Eleventh edition. (Wolters Kluwer Health Adis, 2012), pp. 1–1616.

9. J. L. Flanagan, P. A. Simmons, J. Vehige, M. D. Willcox, Q. Garrett, Role of carnitine in disease. Nutrition & metabolism 7, 30 (2010).

10. M. Almannai, M. Alfadhel, A. W. El-Hattab, Carnitine Inborn Errors of Metabolism. Molecules 24, (2019).

11. T. K. Chen, D. H. Knicely, M. E. Grams, Chronic Kidney Disease Diagnosis and Management: A Review. JAMA 322, 1294–1304 (2019).

12. B. Sharma, D. K. Yadav, L-Carnitine and Chronic Kidney Disease: A Comprehensive Review on Nutrition and Health Perspectives. J Pers Med 13, (2023).

13. X. Yin et al., Genome-wide association studies of metabolites in Finnish men identify disease-relevant loci. Nature communications 13, 1644 (2022).

14. M. C. Costanzo et al., The Type 2 Diabetes Knowledge Portal: An open access genetic resource dedicated to type 2 diabetes and related traits. Cell metabolism 35, 695–710 e696 (2023).

15. P. Dornbos et al., Evaluating human genetic support for hypothesized metabolic disease genes. Cell metabolism 34, 661–666 (2022).

16. G. C. Ferreira, M. C. McKenna, L-Carnitine and Acetyl-L-carnitine Roles and Neuroprotection in Developing Brain. Neurochem Res 42, 1661–1675 (2017).

17. J. S. Yook et al., The SLC25A47 locus controls gluconeogenesis and energy expenditure. Proceedings of the National Academy of Sciences of the United States of America 120, e2216810120 (2023).

18. V. Porcelli, G. Fiermonte, A. Longo, F. Palmieri, The human gene SLC25A29, of solute carrier family 25, encodes a mitochondrial transporter of basic amino acids. The Journal of biological chemistry 289, 13374–13384 (2014).

19. L. Chen et al., Quantitative dynamics of intracellular NMN by genetically encoded biosensor. Biosens Bioelectron 267, 116842 (2025).

20. A. Khan et al., Machine-learning-guided discovery of SLC25A45 as a mediator of mitochondrial methylated amino acid import and carnitine synthesis. Cell metabolism 37, 2220–2232 e2228 (2025).

21. M. M. Dias et al., SLC25A45 is required for mitochondrial uptake of methylated amino acids and de novo carnitine biosynthesis. Molecular cell 85, 4093–4104 e4098 (2025).

22. S. Ravaud et al., Impaired transport of nucleotides in a mitochondrial carrier explains severe human genetic diseases. ACS Chem Biol 7, 1164–1169 (2012).

23. J. Mifsud et al., The substrate specificity of the human ADP/ATP carrier AAC1. Mol Membr Biol 30, 160–168 (2013).

24. J. Tai et al., Hem25p is required for mitochondrial IPP transport in fungi. Nature cell biology 25, 1616–1624 (2023).

25. M. Varadi et al., AlphaFold Protein Structure Database in 2024: providing structure coverage for over 214 million protein sequences. Nucleic acids research 52, D368–D375 (2024).

26. J. J. Ruprecht, E. R. S. Kunji, The SLC25 Mitochondrial Carrier Family: Structure and Mechanism. Trends Biochem Sci 45, 244–258 (2020).

27. A. R. P. Verkerke et al., SLC25A48 controls mitochondrial choline import and metabolism. Cell metabolism 36, 2156–2166 e2159 (2024).

28. S. Patil et al., The membrane transporter SLC25A48 enables transport of choline into human mitochondria. Kidney Int 107, 296–301 (2025).

29. A. Khan et al., Metabolic gene function discovery platform GeneMAP identifies SLC25A48 as necessary for mitochondrial choline import. Nature genetics 56, 1614–1623 (2024).

30. K. Novakova et al., Effect of L-carnitine supplementation on the body carnitine pool, skeletal muscle energy metabolism and physical performance in male vegetarians. Eur J Nutr 55, 207–217 (2016).

31. G. Sveinbjornsson et al., Rare mutations associating with serum creatinine and chronic kidney disease. Human molecular genetics 23, 6935–6943 (2014).

32. C. Tabula Muris et al., Single-cell transcriptomics of 20 mouse organs creates a Tabula Muris. Nature 562, 367–372 (2018).

33. X. Shao, S. Somlo, P. Igarashi, Epithelial-specific Cre/lox recombination in the developing kidney and genitourinary tract. J Am Soc Nephrol 13, 1837–1846 (2002).

34. P. Schonfeld, L. Wojtczak, Short- and medium-chain fatty acids in energy metabolism: the cellular perspective. Journal of lipid research 57, 943–954 (2016).

35. A. S. Pereyra, K. L. McLaughlin, K. A. Buddo, J. M. Ellis, Medium-chain fatty acid oxidation is independent of l-carnitine in liver and kidney but not in heart and skeletal muscle. Am J Physiol Gastrointest Liver Physiol 325, G287–G294 (2023).

36. Q. Liu, Z. Zhou, P. Liu, S. Zhang, Comparative proteomic study of liver lipid droplets and mitochondria in mice housed at different temperatures. FEBS letters 593, 2118–2138 (2019).

37. J. Simcox et al., Global Analysis of Plasma Lipids Identifies Liver-Derived Acylcarnitines as a Fuel Source for Brown Fat Thermogenesis. Cell metabolism 26, 509–522 e506 (2017).

38. S. Gabery, et al., Semaglutide lowers body weight in rodents via distributed neural pathways. JCI Insight 5, (2020).

39. E. Ravussin et al., Tirzepatide did not impact metabolic adaptation in people with obesity, but increased fat oxidation. Cell metabolism, (2025).

40. K. Yokogawa et al., Decreased tissue distribution of L-carnitine in juvenile visceral steatosis mice. The Journal of pharmacology and experimental therapeutics 289, 224–230 (1999).

41. T. Inagaki et al., Endocrine regulation of the fasting response by PPARalpha-mediated induction of fibroblast growth factor 21. Cell metabolism 5, 415–425 (2007).

42. S. A. Kliewer, D. J. Mangelsdorf, A Dozen Years of Discovery: Insights into the Physiology and Pharmacology of FGF21. Cell metabolism 29, 246–253 (2019).

43. M. A. Paquette et al., Thyroid hormone-regulated gene expression in juvenile mouse liver: identification of thyroid response elements using microarray profiling and in silico analyses. BMC Genomics 12, 634 (2011).

44. S. Galland et al., Thyroid hormone controls carnitine status through modifications of gamma-butyrobetaine hydroxylase activity and gene expression. Cellular and molecular life sciences: CMLS 59, 540–545 (2002).

45. M. S. Jansen, G. A. Cook, S. Song, E. A. Park, Thyroid hormone regulates carnitine palmitoyltransferase Ialpha gene expression through elements in the promoter and first intron. The Journal of biological chemistry 275, 34989–34997 (2000).

46. C. M. Kusminski et al., Transforming obesity: The advancement of multi-receptor drugs. Cell 187, 3829–3853 (2024).

47. D. J. Drucker, Prevention of cardiorenal complications in people with type 2 diabetes and obesity. Cell metabolism 36, 338–353 (2024).

48. D. J. Drucker, GLP-1 physiology informs the pharmacotherapy of obesity. Molecular metabolism 57, 101351 (2022).

49. P. Squire, J. Naude, A. Zentner, J. Bittman, N. Khan, Factors associated with weight loss response to GLP-1 analogues for obesity treatment: a retrospective cohort analysis. BMJ Open 15, e089477 (2025).

50. J. Klen, V. Dolzan, Glucagon-like Peptide-1 Receptor Agonists in the Management of Type 2 Diabetes Mellitus and Obesity: The Impact of Pharmacological Properties and Genetic Factors. Int J Mol Sci 23, (2022).

51. M. Javorsky et al., A missense variant in GLP1R gene is associated with the glycaemic response to treatment with gliptins. Diabetes, obesity & metabolism 18, 941–944 (2016).

52. D. A. de Luis, G. Diaz Soto, O. Izaola, E. Romero, Evaluation of weight loss and metabolic changes in diabetic patients treated with liraglutide, effect of RS 6923761 gene variant of glucagon-like peptide 1 receptor. J Diabetes Complications 29, 595–598 (2015).

53. M. Askarpour et al., Beneficial effects of l-carnitine supplementation for weight management in overweight and obese adults: An updated systematic review and dose-response meta-analysis of randomized controlled trials. Pharmacol Res 151, 104554 (2020).

54. M. Pooyandjoo, M. Nouhi, S. Shab-Bidar, K. Djafarian, A. Olyaeemanesh, The effect of (L-)carnitine on weight loss in adults: a systematic review and meta-analysis of randomized controlled trials. Obesity reviews: an official journal of the International Association for the Study of Obesity 17, 970–976 (2016).

55. U. L. Aune, L. Ruiz, S. Kajimura, Isolation and differentiation of stromal vascular cells to beige/brite cells. Journal of visualized experiments: JoVE, (2013).

56. H. Ohno, K. Shinoda, B. M. Spiegelman, S. Kajimura, PPARgamma agonists induce a white-to-brown fat conversion through stabilization of PRDM16 protein. Cell metabolism 15, 395–404 (2012).

57. K. Shinoda et al., Genetic and functional characterization of clonally derived adult human brown adipocytes. Nature medicine 21, 389–394 (2015).

